## Supplementary figures and tables for "The Evolution of Centriole Degradation in Mouse Sperm"

### Supplementary Figures and Legends

**SFig 1-1 – Spermatozoa with one canonical centriole (the proximal centriole) and one atypical centriole (the distal centriole) are present throughout the mammalian tree of life.**

**A**) Example of paddle/eyedrop-shaped sperm head with a centrally inserted neck, barrel-shaped proximal centriole (PC, canonical centriole), and funnel-shaped distal centriole (DC, atypical centriole), as in human, and a sickle-shaped sperm head with a laterally attached neck and no centrioles, as in house mice. Based on Knobil and Neill's Physiology of Reproduction (Plant and Zeleznik, 2014). The acrosome is shown in red, the nucleus is shown in blue, and all other parts of the sperm are shown in green. **B**) A systematic survey of previous transmission electron microscopic studies of Eutherian spermatozoan centrioles.

A

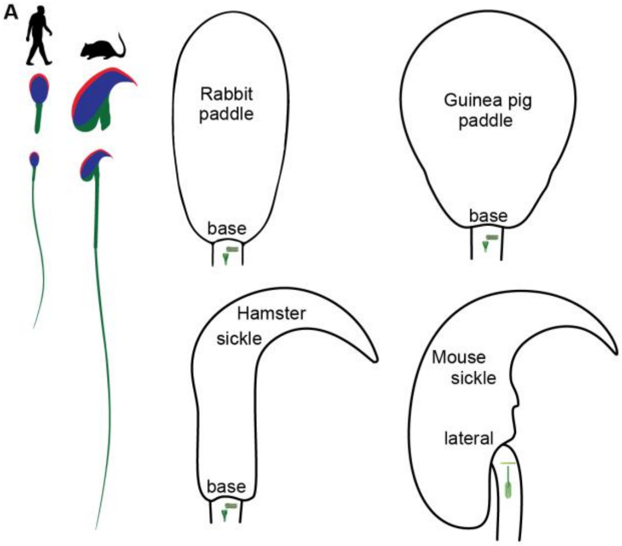


**B Bi) Eutherian**: The four main groups of Eutherian mammalian species (color-coded) have one canonical centriole (the PC) in their spermatozoa. Phylogeny based on (Song et al., 2012).

Phylogeny: Boreoeutheria; Euarchontoglires (Superorder) - Order: Primates
African green monkey (*Cercopithecus aethiops*) (Bedford, 1967)
*Homo sapiens* (Pedersen, 1969) (Garanina et al., 2019)
Phylogeny: Boreoeutheria; Euarchontoglires (Superorder); Glires Order: Rodentia
Siberian chipmunk (*Tamias sibiricus*) (Lee and Park, 2011)
Chinchilla (Healey and Weir, 1970)
Guinea Pig (Caviidae) (Fawcett, 1965)

Phylogeny: Boreoeutheria; Laurasiatheria (Superorder) - Order: Carnivora
Cat (Schmehl and Graham, 1989) (Sato and Oura, 1984)
Phylogeny: Boreoeutheria; Laurasiatheria (Superorder) - Order: Chiroptera
Bat (Bernard and Hodgson, 1988)
Phylogeny: Boreoeutheria; Laurasiatheria (Superorder) - Order: Eulipotyphla
Lesser white-toothed shrew (*Crocidura suaveolens*) (Soon-Jeong et al., 2006)
Phylogeny: Boreoeutheria; Laurasiatheria (Superorder) - Order: Artiodactyla
Boar (Nicander and Bane, 1962)
Bovine (Ounjai et al., 2012)
Camel ` (Tingari, 1991)
Phylogeny: Boreoeutheria; Laurasiatheria (Superorder) - Order: Perissodactyla
Horse (Leung et al., 2021)

Phylogeny: Afrotheria; Order: Proboscidea
Asiatic elephant (*Elephas maximus*) (Heath et al., 1983)
Phylogeny: Afrotheria Order: Macroscelidea
Macroscelidea (elephant shrews) (Woodall and FitzGibbon, 1995)

Phylogeny: Xenarthra
Six-banded armadillo (*Euphractus sexcinctus*) (Sousa et al., 2013)

**Bii) Monotreme**: The structure of the sperm neck is simpler in Monotreme Mammals than in Eutherian Mammals: While a PC is found near the nucleus, and the mitochondria reach the nucleus, neither striated columns nor a canonical DC are observed (Carrick and Hughes, 1982).

**Biii) Marsupial**: In Marsupials, the DC is not observed or persists in a modified form (Lloyd et al., 2002; Temple-Smith, 1994). The PC (aka transverse centriole) is observed in some species (Lloyd et al., 2002), having been reported in the mature spermatozoa of *Perameles nasuta*, *Macrotis lagotis*, several dasyurids, the petaurid genus Petaurus, and the peramelid *Isoodon macrourus* ((Sapsford et al., 1969) and (Johnston et al., 1995) in (Lloyd et al., 2002)). A remnant PC is present in *Hypsiprymnodon moschatus* (Lloyd et al., 2002). No PC is detected in *Trichosurus vulpecula* and most macropods ((Harding, 1979) in (Lloyd et al., 2002)).

**SFig 1B References:**


Bedford, J.M. 1967. Observations on the fine structure of spermatozoa of the bush baby (Galago senegalensis), the African green monkey (Cercopithecus aethiops) and man. *The American journal of anatomy*. 121:443-459.

Bernard, R.T., and A.N. Hodgson. 1988. Fine structure of the neck of epididymal spermatozoa of Schreiber's long-fingered bat (Chiroptera: Mammalia). *Gamete Res*. 21:41-50.

Carrick, F.N., and R.L. Hughes. 1982. Aspects of the Structure and Development of Monotreme Spermatozoa and Their Relevance to the Evolution of Mammalian Sperm Morphology. *Cell and Tissue Research*. 222:127-141.

Fawcett, D.W. 1965. The anatomy of the mammalian spermatozoon with particular reference to the guinea pig. *Z Zellforsch Mikrosk Anat*. 67:279-296.

Garanina, A.S., I.B. Alieva, E.E. Bragina, E. Blanchard, B. Arbeille, F. Guerif, S. Uzbekova, and R.E. Uzbekov. 2019. The Centriolar Adjunct(-)Appearance and Disassembly in Spermiogenesis and the Potential Impact on Fertility. *Cells*. 8:180.

Harding, H.R. 1979. Reproduction in male marsupials. A critique with additional observation on sperm development and structure, Elibrary..

Healey, P., and B.J. Weir. 1970. Changes in the ultrastructure of chinchilla spermatozoa in different diluents. *J Reprod Fertil*. 21:191-193.

Heath, E., R. Jeyendran, and E. Graham. 1983. Ultrastructure of Spermatozoa of the Asiatic Elephnat (Elephas maximus). *Anatomia, Histologia, Embryologia*. 12:245-252.

Johnston, S., L. Daddow, and F. Carrick. 1995. Ultrastructural and light microscopic observations of mature epididymal spermatozoa and sperm maturation of the greater bilby Macrotis lagotis (Metatheria, Mammalia). *Advances in Spermatozoal Phylogeny and Taxonomy*. 166:397-407.

Lee, J.-H., and K.-R. Park. 2011. Fine Structure of Sperm in the Korea Squirrel, Tamias sibiricus. *Applied Microscopy*. 41:99-107.

Leung, M.R., M.C. Roelofs, R.T. Ravi, P. Maitan, H. Henning, M. Zhang, E.G. Bromfield, S.C. Howes, B.M. Gadella, H. Bloomfield-Gadelha, and T. Zeev-Ben-Mordehai. 2021. The multi-scale architecture of mammalian sperm flagella and implications for ciliary motility. *EMBO J*. 40:e107410.

Lloyd, S., F. Carrick, and L. Hall. 2002. Ultrastructure of the mature spermatozoon of the musky rat-kangaroo, Hypsiprymnodon moschatus (Potoroidae: Marsupialia). *Acta Zoologica*. 83:167-174.

Nicander, L., and A. Bane. 1962. Fine structure of boar spermatozoa. *Z Zellforsch Mikrosk Anat*. 57:390-405.

Ounjai, P., K.D. Kim, P.V. Lishko, and K.H. Downing. 2012. Three-dimensional structure of the bovine sperm connecting piece revealed by electron cryotomography. *Biology of reproduction*. 87:73.

Pedersen, H. 1969. Ultrastructure of the ejaculated human sperm. *Z Zellforsch Mikrosk Anat*. 94:542-554.

Sapsford, C., C.A. Rae, and K. Cleland. 1969. Ultrastructual studies on maturing spermatids and on sertoli cells in the bandicoot Perameles nasuts Geoffroy (Marsupialia). *Australian Journal of Zoology*. 17:195-292.

Sato, N., and C. Oura. 1984. The fine structure of the neck region of cat spermatozoa. *Okajimas folia anatomica Japonica*. 61:267-285.

Schmehl, M., and E. Graham. 1989. Ultrastructure of the domestic tom cat (Felisdomestica) and tiger (Pantheratigrisaltaica) spermatozoa. *Theriogenology*. 31:861-874.

Song, S., L. Liu, S.V. Edwards, and S.Y. Wu. 2012. Resolving conflict in eutherian mammal phylogeny using phylogenomics and the multispecies coalescent model. *Proceedings of the National Academy of Sciences of the United States of America*. 109:14942-14947.

Soon-Jeong, J., P. Joo-Cheol, K. Heung-Joong, S.B. Chun, Y. Myung-Hee, L. Do-Seon, and J. Moon-Jin. 2006. Comparative fine structure of the epididymal spermatozoa from three Korean shrews with considerations on their phylogenetic relationships. *Biocell*. 30:279-286.

Sousa, P.C., E.A. Santos, J.A. Bezerra, G.L. Lima, T.S. Castelo, J.D. Fontenele-Neto, and A.R. Silva. 2013. Morphology, morphometry and ultrastructure of captive six-banded armadillo (Euphractus sexcinctus) sperm. *Animal reproduction science*. 140:279-285.

Temple-Smith, P.D. 1994. Comparative structure and function of marsupial spermatozoa. *Reprod Fertil Dev*. 6:421-435.

Tingari, M. 1991. Studies on camel semen. III. Ultrastructure of the spermatozoon. *Animal reproduction science*. 26:333-344.

Woodall, P.F., and C. FitzGibbon. 1995. Ultrastructure of Spermatozoa of the Yellow-rumped Elephant Shrew Rhynchocyon chrysopygus (Mammalia: Macroscelidea) and the Phylogeny of Elephant Shrews. *Acta Zoologica*. 76:19-23.

**SFig 1-2 – Survey of original literature on rodent spermatozoan ultrastructure**

This figure is a large data set. See supplementary files**.**

**STable 2-1 – Identity Ratio range of sperm centrosomal proteins**

This table is a large data set. See supplementary file**.**

**STable 2-2 – In rodents, natural selective pressure on FAM161A is stronger than on other rod proteins and is strongest in Muridae.**

| **Gene** | *FAM161A* (Muridae) | *FAM161A* (Cricetidae) | *FAM161A* (Other Myomorpha) | *FAM161A* (Rodents) | *FAM161B* (Rodents) | *POC1b* (Rodents) | *POC5* (Rodents) | *CETN1* (Rodents) | *WDR90* (Rodents) |
|---|---|---|---|---|---|---|---|---|---|
| ***ω (dN/dS)*** | 0.62960 | 0.57030 | 0.48616 | 0.57592 | 0.33602 | 0.19055 | 0.26702 | 0.03423 | 0.26510 |

**STable 2-3 – Codon-wide selection analysis to identify sites under positive and negative selection**

Selection analysis identified more positively (Green font) and negatively (red font) selected sites in rodents than in primates, Carnivora, and ungulates. The number of codons used in each lineage is indicated in parentheses**.** The number of selected sites and the site numbers along the primary structure are indicated for each method and mammalian order. MEME (Murrell et al., 2012) is a generalization of FEL (Kosakovsky Pond and Frost, 2005). The initial analytical phases of the two approaches are identical; however, FEL assumes that the same dN/dS (ω) ratio applies to all branches, while MEME models variable dN/dS (ω) values across lineages at an individual site.

| **Selection Analysis** | **Ungulates** 722 codons | **Carnivora**722 c odons | **Primates** 716 codons | **Rodents** 714 codons |
|---|---|---|---|---|
| **MEME:** Episodic Diversifying Selection (P< 0.05) | **9 Sites**: 46, 47, 49, 60, 62, 67, 84, 312, 440, 505 | **7 sites**: (35, 49, 60, 65, 451, 476, 518, 688 | **10 Sites**: 44, 53, 136, 186, 268, 302, 348, 633, 634, 695 | **53 Sites**: 5, 10, 11, 15, 18, 22, 23, 44, 56, 59, 60, 61, 62, 65, 67, 68, 71, 113, 134, 135, 174, 177, 230, 237, 254, 325,378, 422, 427, 442, 451, 453, 467, 479, 633, 634, 635, 636, 639, 657, 680, 682, 687, 690, 713 |
| **FEL:** Pervasive Diversifying Selection (P< 0.05) | **0 Sites** | **2 Sites**: (27, 65) | **2 Sites**: 54, 268 | **14 Sites:**10, 22, 23, 62, 67, 230, 363, 392, 427, 442, 451, 473, 512, 656 |
| **PAML** (M8 BEB, No *= PP=90, *PP >95, **PP>95) | **5 Sites:** 13L, 46L, 61A, 61R*, 511A* | **17 Sites:** 5H, 17T, 27V, 35P, 36L*, 65R*, 85G, 172C*, 219Q, 315Y*, 420G, 429C, 450H, 476T*, 498Y*, 589M*, 603Q* | **6 Sites:** 38A, 171V, 213R*, 306R, 391H*, 406C**) | **42 Sites:** 4P, 22 I**, 45A, 46A**, 48M, 52E**, 53Q**, 55K, 56V, 65G*, 67H*, 71G*, 78F, 81T, 129F, 130I, 152L*, 184T*, 192V, 213*, 221A, 230S, 264*, 270R*, 288S, 289C*, 345F, 399S, 403C**, 405R*, 406F, 438W**, 442P*, 448F*, 451C*, 455C*,467S, 473L*, 496R*, 511E*, 512C*, 656F |
| **FEL:** Purifying Selection (P< 0.05) | **34 sites:** 3, 50, 51, 54, 55, 56, 107, 135, 146, 159, 166, 232, 235, 277, 291, 294, 295, 298, 310, 323, 330, 331, 359, 361, 371, 390, 463, 465, 523, 526, 530, 567, 583, 603 | **38 Sites** 37,38,45,78,91, 130,159,163,18 2,203,204,212,1 49,252,264,273, 280,292,296,32 8,330,373,398,4 02,412,436,447, 461,469,479,52 2,526,530,535, 555,617,634,71 4 | **37 Sites** 8,12,34,35,58,6 2,71,76,83,94,1 53,160,178,198 ,208,211,239,2 46,262,273,326 ,331,369,404,4 12,444,452,453 ,486,517,558,5 64,585,586,637 ,659,664 | **100 Sites:** 2, 8, 9, 51, 86, 90, 93, 97, 98, 101, 103, 105, 111, 121, 138, 140, 143, 155, 156, 158, 165, 170, 185, 194, 198, 205,210,236,240,241,244,245,246,258, 274,285,287,292,293,295,300,303,304, 305,306,318,321,323,324,326,327,329, 330,331,335,337,338,357,358,371,379, 389,425,464,484,499,514,518,522,523, 534,525,541,548,569,570,575,591,595, 598,605,606,610,611,612,615,616,620, 622,624,625,626,637,638,654,665,674, 679,695,707 |
| **FUBAR:** Pervasive Purifying Selection (P< 90) | **28 Sites:** 3, 50, 51, 54, 55, 56, 60, 107, 135, 146, 159, 166, 232, 235, 277, 294, 310, 325, 330, 359, 390, 465, 523, 526, 530, 567, 583, 603 | **31 Sites:** 37, 38, 45, 78, 91, 130, 159, 182, 203, 212, 253, 264, 273, 292, 296, 328, 330, 398, 412, 447, 461, 469, 479, 522, 526, 530, 535, 555, 617, 634 | **19 Sites:** 34, 58, 62, 76, 178, 208, 239, 262, 273, 326, 404, 412, 444, 452, 486, 517, 586, 659, 664 | **107 Sites:** 1, 2, 9, 86, 90, 93, 94, 98, 101, 103, 105, 109, 115, 138, 140, 143, 155, 156, 158, 165, 170, 185, 194, 198, 201, 205, 209, 210, 236, 238, 240, 241, 243, 244, 246, 258, 274, 276, 280, 281, 282, 283, 285, 287, 293, 295, 304, 306, 318, 323, 324, 326, 329, 330, 331, 335, 337, 338, 354, 356, 358, 364, 371, 465, 470, 481, 514, 518, 523, 524, 525, 527, 533, 537, 548, 563, 565, 570, 575, 585, 591, 592, 595, 598, 605, 606, 610, 612, 613, 614, 615, 616, 620, 621, 622, 624, 625, 626, 637, 638, 650, 663, 665, 674, 693, 695, 707 |

**STable 2-4 – Number of negatively selected (NSS) and positively selected (PSS) sites with P<0.05 using FEL and FUBAR analysis.**

| | | Muridae | Cricetidae | Other rodents |
|---|---|---|---|---|
| FEL (P<0.05) | Negatively selected sites (NSS) | 4 | 87 | 48 |
| | Positively selected sites (PSS) | 14 | 14 | 15 |
| FUBAR (PP >90) | Negatively selected sites (NSS) | 7 | 14 | 28 |
| | Positively selected sites (PSS) | 0 | 3 | 0 |

**SFig 3-1 – House mice express a Type 3 (555-aa) FAM161A isoform in the testes.**

**A**) Various annotated FAM161A isoforms found in house mouse testes, based on transcriptome sequencing performed by (Green et al., 2018). **B–C**) FAM161A isoform types 1–3 with exons numbered and approximate locations of forward, reverse (filled, colored triangles), and sequencing primers (open, colored triangles) chosen for PCR with reverse-transcribed mRNA isolated from either house mouse testis or eye tissue (**B**); and electrophoresis gel with expected sizes (in base pairs) of bands produced using the indicated primers (**C**). -, no band was expected for that reaction. A cDNA library was constructed using a 3' RACE kit (Roche). The First PCR was performed using the cDNA library, the product of which was used to perform the Second PCR. Sequencing was performed on the Second PCR product following gel extraction. MW, molecular weight. Results were consistent across three independent experiments. Uncropped gel images are provided in **SFig 3-3. D**) The NCBI house mouse testis EST database showed that eight FAM161A transcripts have the unique C-terminus sequence characteristic of type 3.

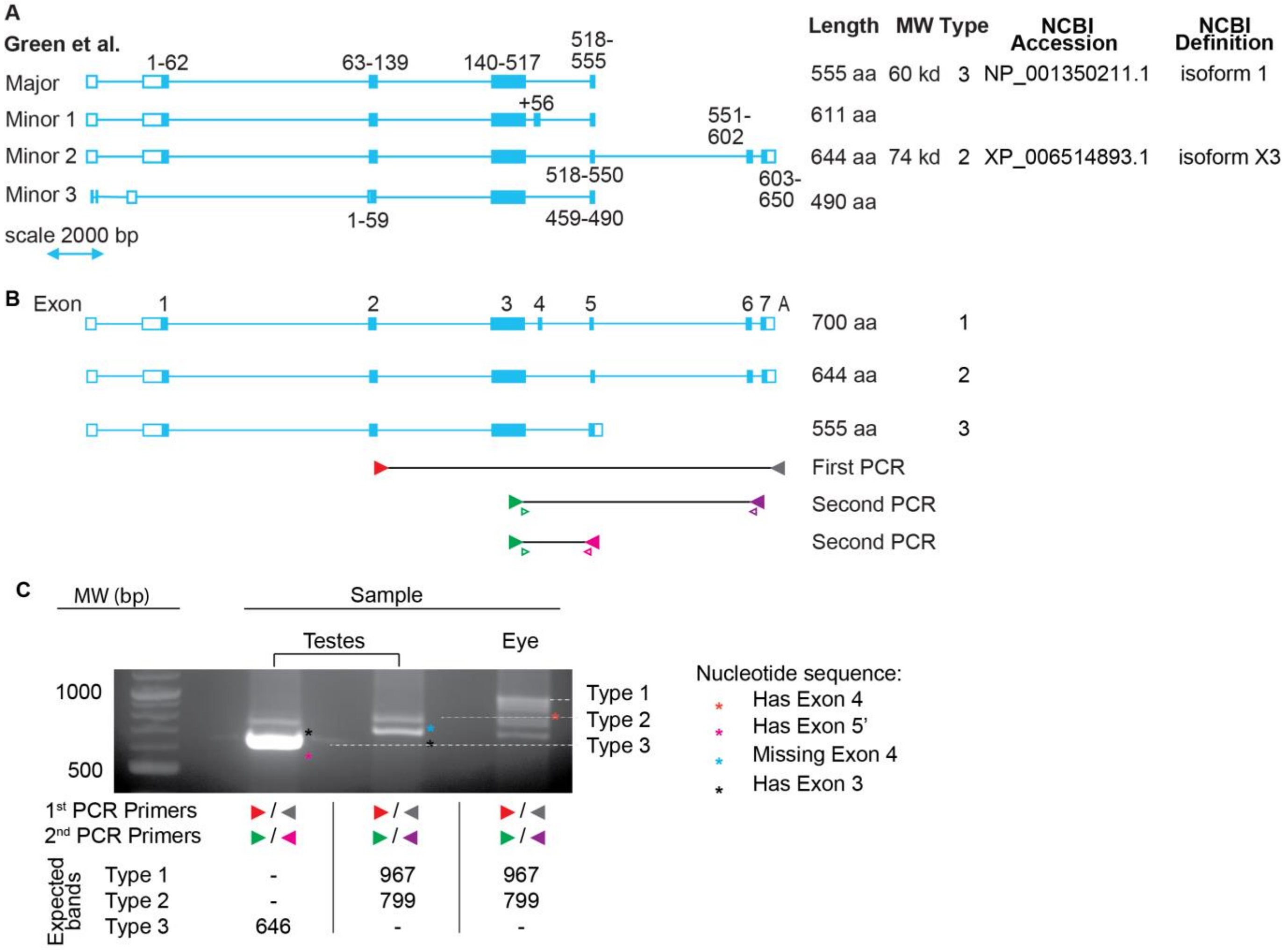


**D** The NCBI mouse testis EST database shows that eight FAM161A transcripts have the unique, C-terminus sequence characteristic of type 3:

AV283297.2, AI613936.1, AV262776.1, AA492939.1, BB015036.1, AV257583.1, AV283863.1, AA064510.1.

**SFig 3-2 – Uncropped images of Western blots**

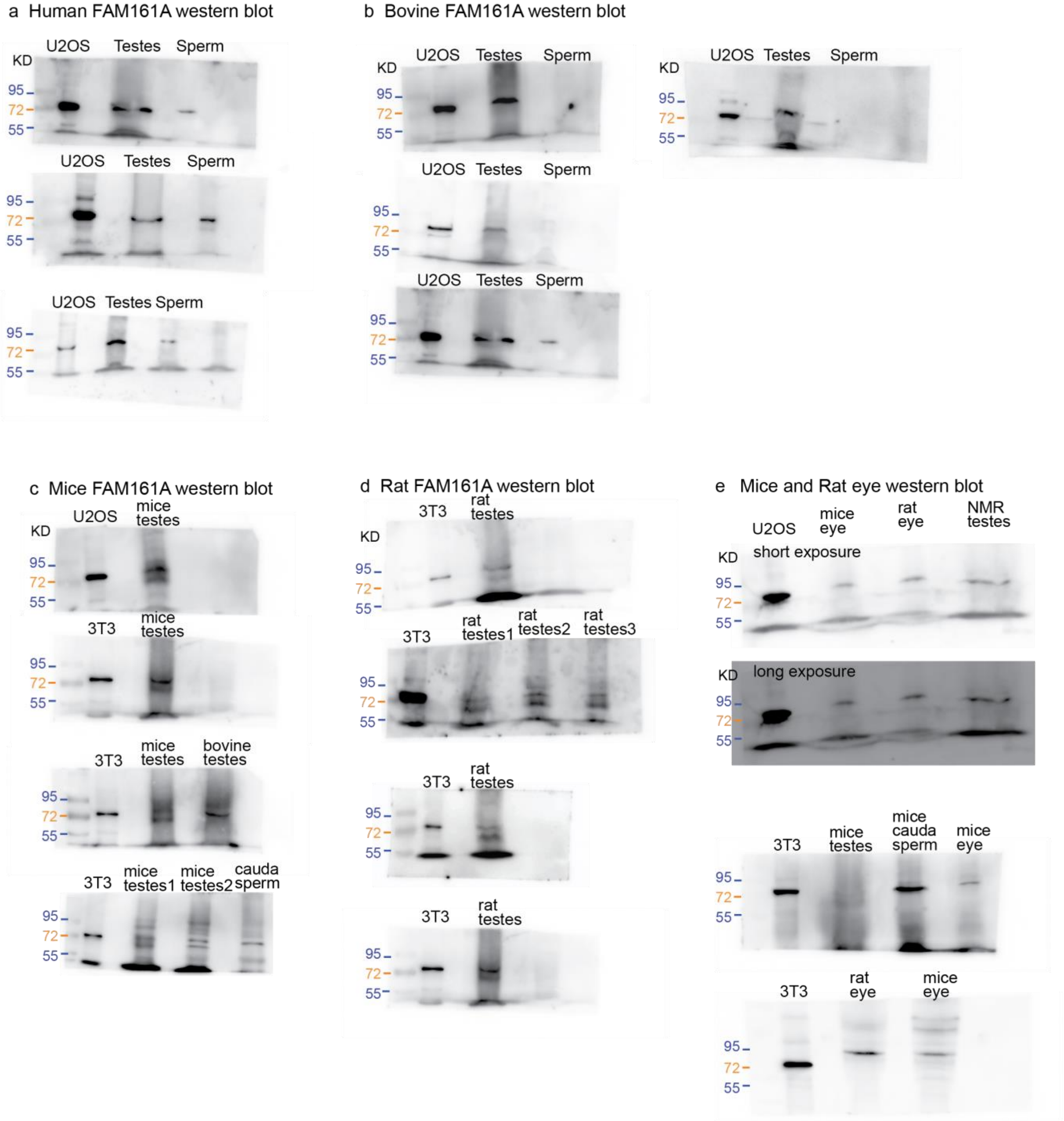

**SFig 3-3 – House mice express a Type 3 (555-aa) FAM161A isoform in the testes.**

**A–C**) Uncropped electrophoresis gels with expected sizes (in base pairs) of bands produced from each tissue type, produced by repeating the experiment described in **SFig 3-1** and captured using a c500 Bioanalytical Imaging System (Azure Biosystems, Inc.) (**A and B**) or a DIGI DOC-IT High Performance Ultraviolet Transilluminator (UVP) (**C**). -, no band was expected for that reaction. MW, molecular weight.

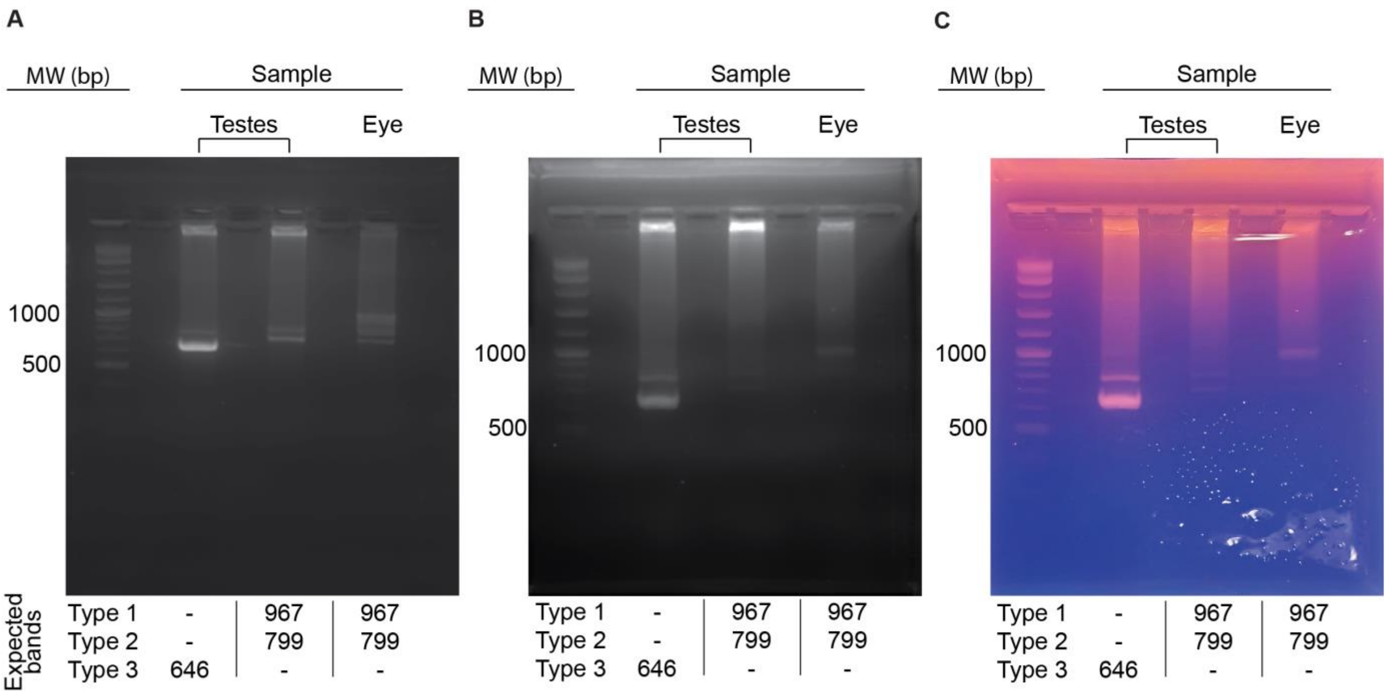

**SFig 4-1 – House mouse and human FAM161A isoforms can localize to canonical centrioles in U2OS cells.**

**A and C**) Human FAM161A (hFAM161A) type 2, house mouse FAM161A (mFAM161A) type 2, and mFAM161A type 3 were localized to the centrosome based on pericentrin (**A**) and γ-tubulin (**C**) labeling in U2OS cells. Human FAM161A type 2 and mFAM161A type 2 also exhibit pericentrosomal localization, probably because these isoforms colocalize with microtubules emanating from the centrosome. **B and D**) Quantification showing the percentage of centrioles exhibiting centrosomal, pericentrosomal, and non-centrosomal localization of hFAM161A type 2, mFAM161A type 2, and mFAM161A type 3. All experiments were repeated three times with consistent results.

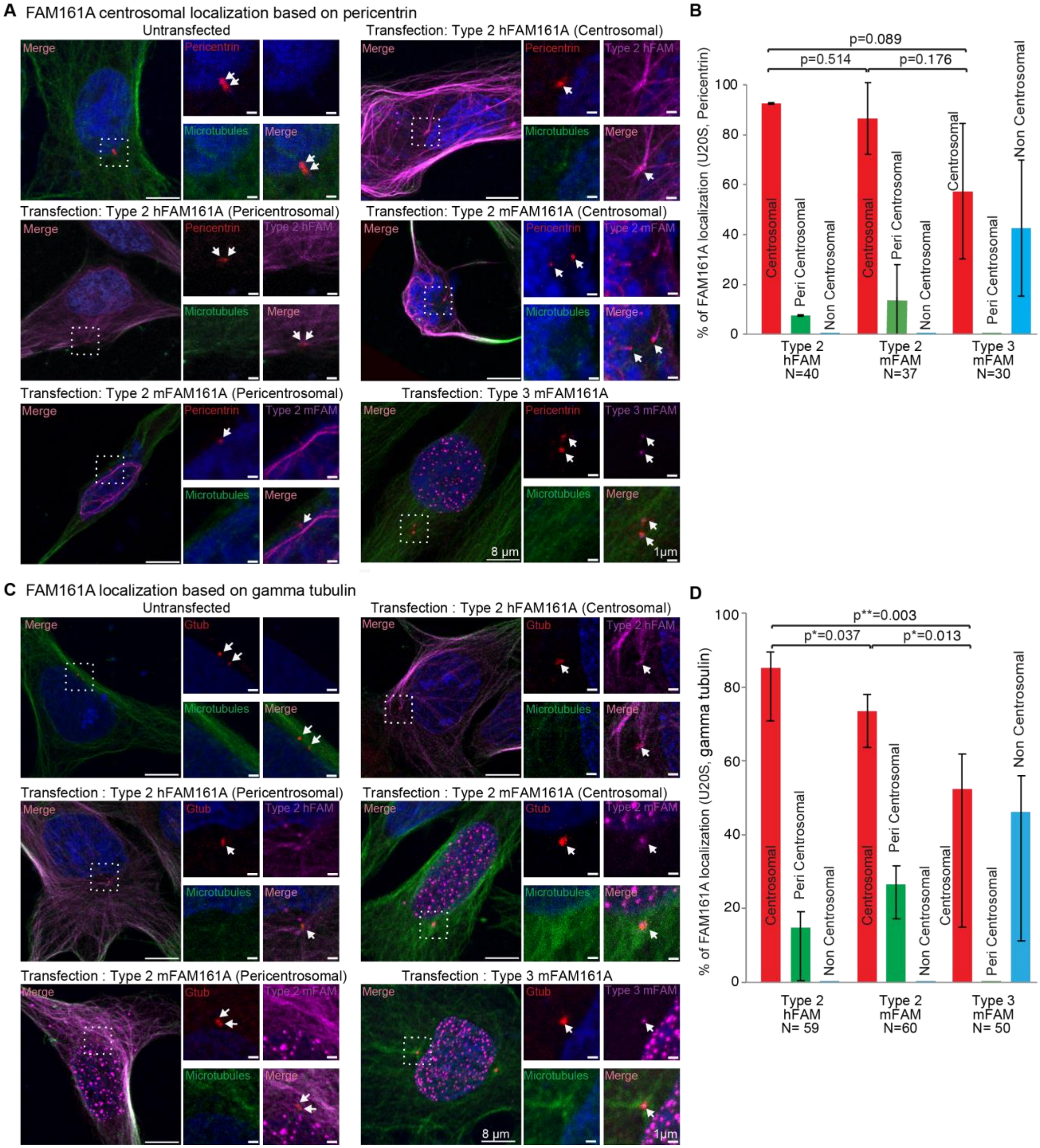

**SFig 4-2 – House mouse and human FAM161A isoforms can localize to canonical centrioles in 3T3 cells.**

**A and C**) Human FAM161A (hFAM161A) type 2, house mouse FAM161A (mFAM161A) type 2, and mFAM161A type 3 were localized to the centrosome based on pericentrin (**A**) and γ-tubulin (**C**) labeling in 3T3 cells. Human FAM161A type 2 and mFAM161A type 2 also exhibit pericentrosomal localization, probably because these isoforms colocalize with microtubules emanating from the centrosome. **B and D**) Quantification showing the percentage of centrioles exhibiting centrosomal, pericentrosomal, and non-centrosomal localization of hFAM161A type 2, mFAM161A type 2, and mFAM161A type 3. All experiments were repeated three times with consistent results.

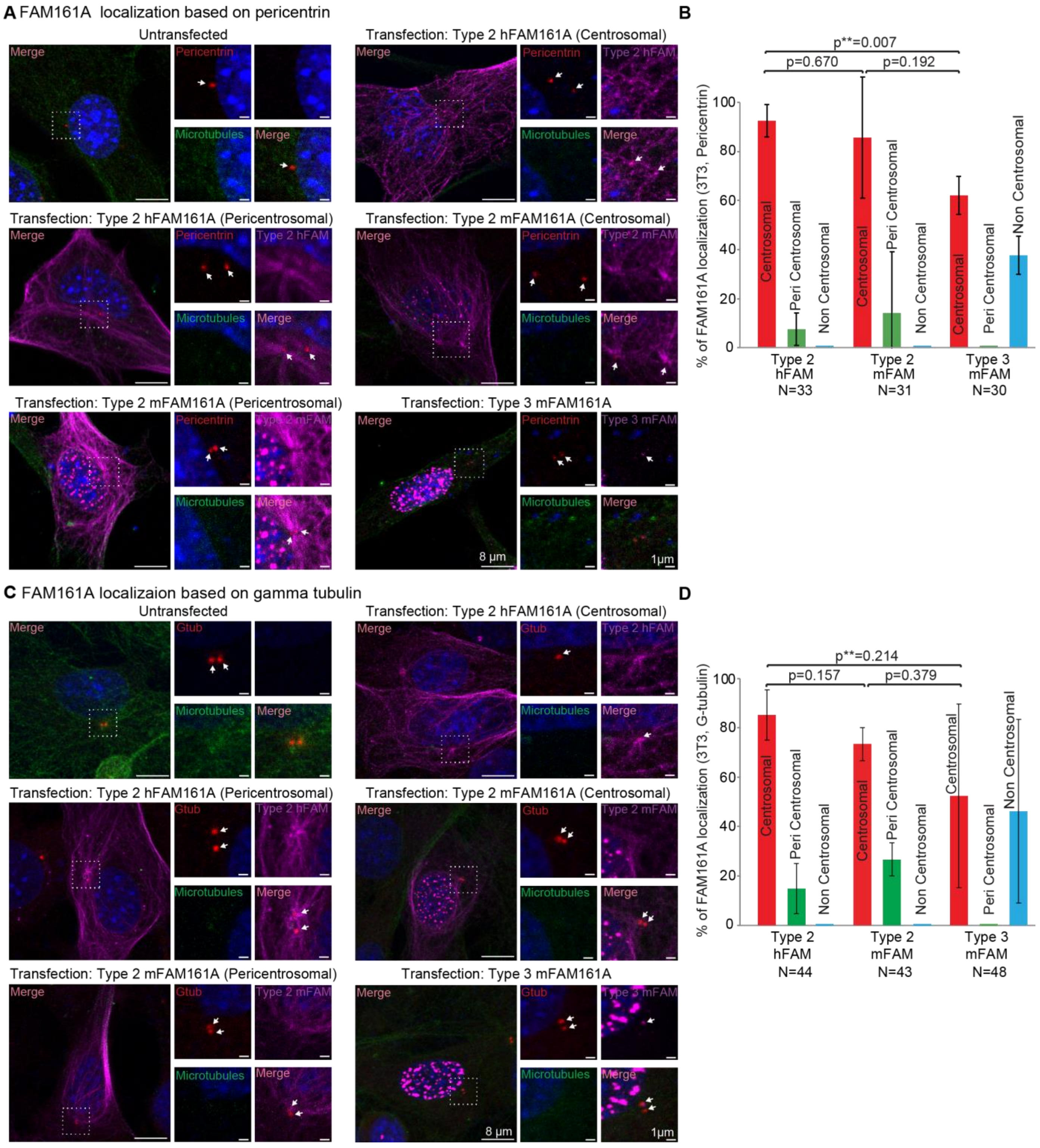
A FAM161A localization based on pericentrin
Untransfected
Transfection: Type 2 hFAM161A (Centrosomal)
Transfection: Type 2 hFAM161A (Pericentrosomal)
Transfection: Type 2 mFAM161A (Centrosomal)
Transfection: Type 2 mFAM161A (Pericentrosomal)
Transfection: Type 3 mFAM161A
Merge
Pericentrin
Microtubules
Type 2 hFAM
Type 2 mFAM
Type 3 mFAM
8 μm
1μm
B
p**=0.007
p=0.670
p=0.192
% of FAM161A localization (3T3, Pericentrin)
Centrosomal
Peri Centrosomal
Non Centrosomal
Type 2 hFAM N=33
Type 2 mFAM N=31
Type 3 mFAM N=30
C FAM161A localizaion based on gamma tubulin
Untransfected
Transfection: Type 2 hFAM161A (Centrosomal)
Transfection: Type 2 hFAM161A (Pericentrosomal)
Transfection: Type 2 mFAM161A (Centrosomal)
Transfection: Type 2 mFAM161A (Pericentrosomal)
Transfection: Type 3 mFAM161A
Merge
Gtub
Microtubules
Type 2 hFAM
Type 2 mFAM
Type 3 mFAM
8 μm
1μm
D
p**=0.214
p=0.157
p=0.379
% of FAM161A localization (3T3, G-tubulin)
Centrosomal
Peri Centrosomal
Non Centrosomal
Type 2 hFAM N=44
Type 2 mFAM N=43
Type 3 mFAM N=48

**SFig 4-3 – The mechanism of rod protein interaction**

**A**) Expression of human FAM161A (hFAM161A) type 2, human POC1B (hPOC1B), and human POC5 (hPOC5) and quantification of their colocalization with tubulin in U2OS cells. **B**) Expression of various fragments of hFAM161A type 2 with full-length hPOC1B and quantification of FAM161A colocalization with hPOC1B. **C**) Expression of various fragments of hFAM161A type 2 with hPOC5 and quantification of FAM161A colocalization with hPOC5. Scale bars are 10 µm. **D**) Yeast two-hybrid analysis to map the hPOC5-interacting domain of hFAM161A. BD, binding domain; AD, activation domain; DDO, double dropout (-Leu, -Trp); TDO, triple dropout (-Leu, -Trp, -His); QDO, quadruple dropout (-Leu, -Trp, -His, -Ade). **E**) Yeast two-hybrid analysis to map the domain mediating the interaction between hPOC1B and hPOC5. **F**) Model depicting the mechanism of interaction between rod proteins. WD, WD40 repeat domains; CBR, centrin-binding region; EF-hand, a calcium-binding motif. CC, coiled-coil domain; n, number of cells. Data shown are the representative images and quantification from three independent experiments.

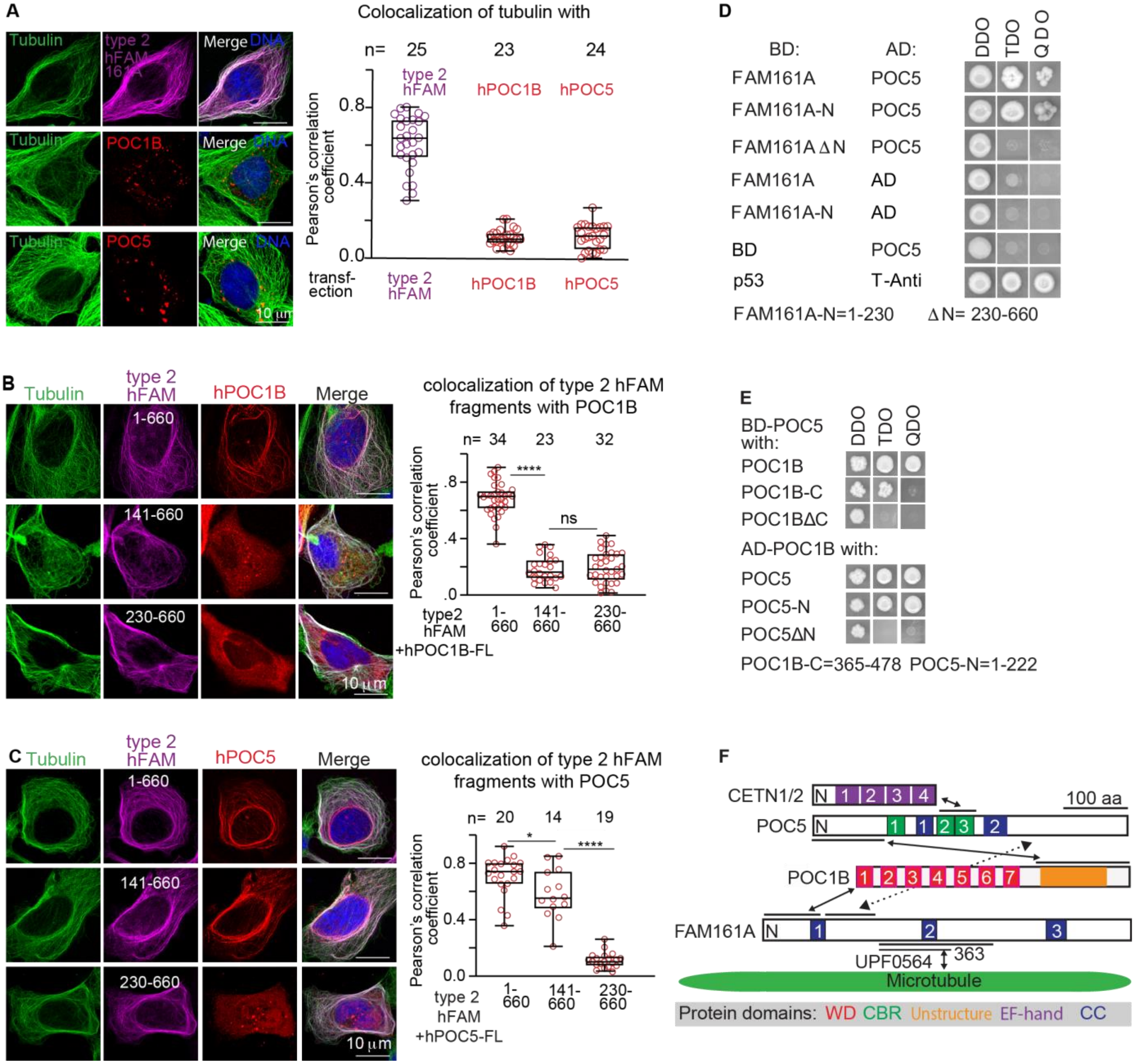

**SFig 4-4 – FAM161A isoforms exhibit different localization patterns in 3T3 cells.**

**A**) Expression analysis of human FAM161A (hFAM161A) type 2 (second panel from left), house mouse FAM161A (mFAM161A) type 2 (middle four panels), mFAM161A type 3 (right panel) in 3T3 cells. **B**) Quantification showing the percentage of cells exhibiting each of the various expression patterns observed during expression analysis of hFAM161A type 2, mFAM161A type 2, and mFAM161A type 3. C, "Cytoplasmic"; I, "Intranuclear"; I+C, "Intranuclear + Cytoplasmic"; P, "Perinuclear". The experiment was repeated three times with consistent results.

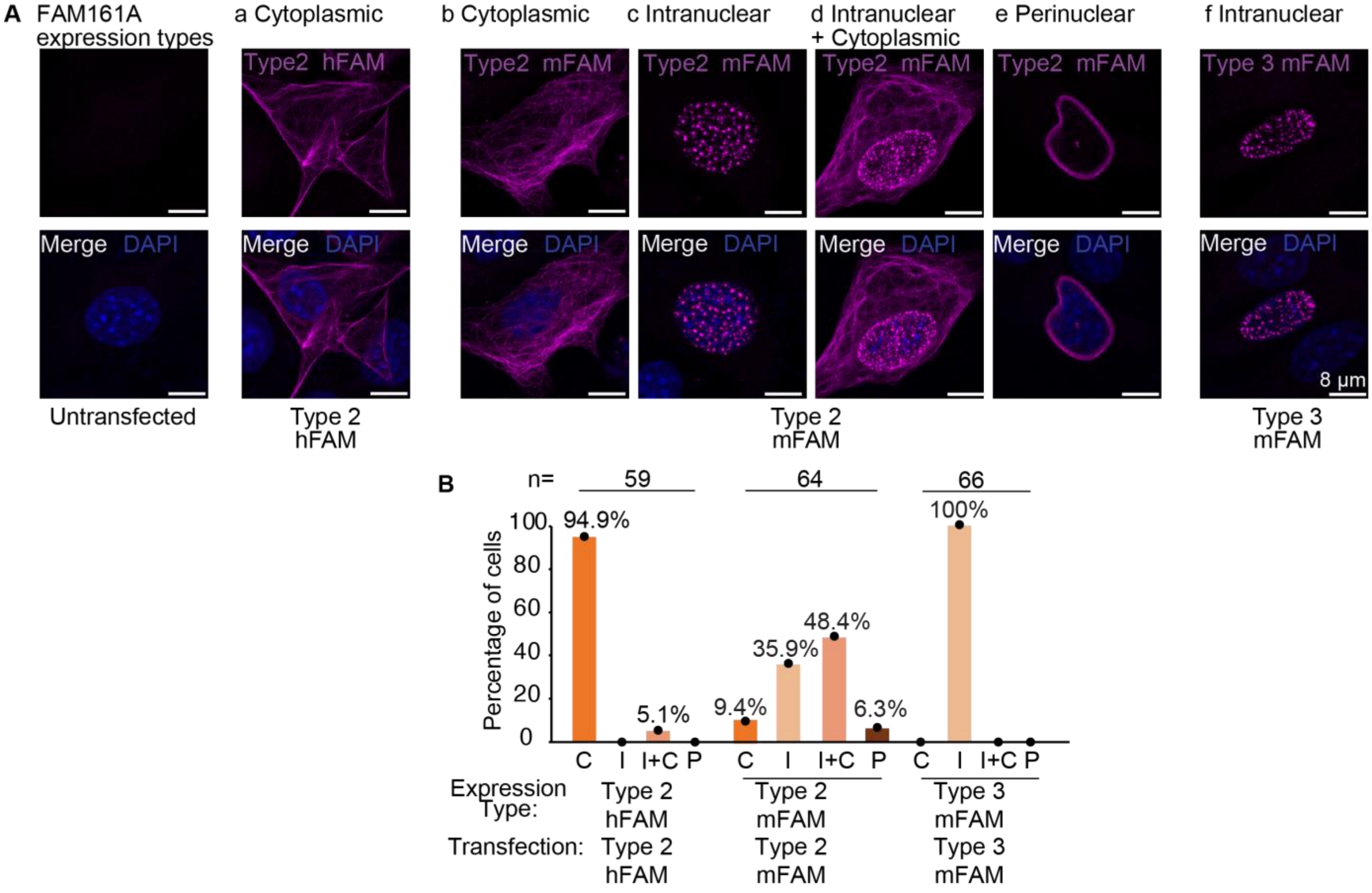

**SFig 4-5 – FAM161A sequence alignment**

Alignment of the amino acid sequences of human FAM161A type 1 (Hs1), human FAM161A type 2 (Hs2), mouse FAM161A type 1 (Mm1), mouse FAM161A type 2 (Mm2), and mouse FAM161A type 3 (Mm3). The various protein domains are marked using color coding.

```
Hs1   MATSHRVAKLVASSLQTPVNPITGARVAQYEREDPLKALAAAEAILEDEEEE--KVAQPA 58
Hs2   MATSHRVAKLVASSLQTPVNPITGARVAQYEREDPLKALAAAEAILEDEEEE--KVAQPA 58
Mm1   MASPHQEARLAAASLLLSANPITGARVAQYESEDSLESLVAAGLAAMEEEEQKKVWQMR 60   POC1B Binding segment
Mm2   MASPHQEARLAAASLLLSANPITGARVAQYESEDSLESLVAAGLAAMEEEEQKKVWQMR 60
Mm3   MASPHQEARLAAASLLLSANPITGARVAQYESEDSLESLVAAGLAAMEEEEQKKVWQMR 60
      **: *: *:*.*:**   .************ ** *::*.**       :**::** *

Hs1   GASADLNTSFSGVDEHAPISYEDFVNFPDIHHSNEEYFKKVEELKAAHIETMAKLEKMYQ 118
Hs2   GASADLNTSFSGVDEHAPISYEDFVNFPDIHHSNEEYFKKVEELKAAHIETMAKLEKMYQ 118
Mm1   -ASAGCH-NFSGKDE--EVAFESTMNFSDIYHSDEEYFRKLKDLKAVHEETMSKLEKMYQ 116
Mm2   -ASAGCH-NFSGKDE--EVAFESTMNFSDIYHSDEEYFRKLKDLKAVHEETMSKLEKMYQ 116
Mm3   -ASAGCH-NFSGKDE--EVAFESTMNFSDIYHSDEEYFRKLKDLKAVHEETMSKLEKMYQ 116
      .  :    .*** **   :::*. :** **:**:****:*:::***.* ***:*******

Hs1   DKLHLKEVQPVVIREDSLSDSSRSVSEKNSYHPVSLMTSFSEPDLGQSSSLYVSSSEEEL 178
Hs2   DKLHLKEVQPVVIREDSLSDSSRSVSEKNSYHPVSLMTSFSEPDLGQSSSLYVSSSEEEL 178   POC5 Binding segment
Mm1   DKLNIKDIQAGFIRDGISDSSSSSASEKSCSHPALSVTSLSEPDLDGSSSLST-TTDEGL 175
Mm2   DKLNIKDIQAGFIRDGISDSSSSSASEKSCSHPALSVTSLSEPDLDGSSSLST-TTDEGL 175
Mm3   DKLNIKDIQAGFIRDGISDSSSSSASEKSCSHPALSVTSLSEPDLDGSSSLST-TTDEGL 175
      ***::*::*  .**:.  ..** *.***.. **.  :**:*****. **** . :::* *

Hs1   PNLEKEYPRKNRMMTYAKELINNMWTDFCVEDYIRCKDTGFHAAEKRRKKRKEWVPTITV 238
Hs2   PNLEKEYPRKNRMMTYAKELINNMWTDFCVEDYIRCKDTGFHAAEKRRKKRKEWVPTITV 238
Mm1   PDLEEKTPGESSAMVHAQELINNMWNDFSVEDYIQYDSDS-RTAKKKRKKAKSLTPKITV 234
Mm2   PDLEEKTPGESSAMVHAQELINNMWNDFSVEDYIQYDSDS-RTAKKKRKKAKSLTPKITV 234
Mm3   PDLEEKTPGESSAMVHAQELINNMWNDFSVEDYIQYDSDS-RTAKKKRKKAKSLTPKITV 234
      *:**:: * :.  *.:*:*******.**.*****: .. . ::*:*:*** *. .*.***

Hs1   PEPFQMMIREQKKKEESMKSKSDIEMVHKALKKQEEDPEYKKKFRANPVPASVFLPLYHD 298
Hs2   PEPFQMMIREQKKKEESMKSKSDIEMVHKALKKQEEDPEYKKKFRANPVPASVFLPLYHD 298   dimerization domain
Mm1   PVPFEMTVREQNRREKALSARSDLE--TKLLKRDEDDAECKKKFRANPVPSCVLLPLYED 292
Mm2   PVPFEMTVREQNRREKALSARSDLE--TKLLKRDEDDAECKKKFRANPVPSCVLLPLYED 292
Mm3   PVPFEMTVREQNRREKALSARSDLE--TKLLKRDEDDAECKKKFRANPVPSCVLLPLYED 292
      * **:* :***:::*:::.::**:*   * **::*:* * **********:.*:****.*

Hs1   LVKQKEERRRSLKEKSKEALLASQKPFKFIAREEQKRAAREKQLRDFLKYKKKTNRFKAR 358
Hs2   LVKQKEERRRSLKEKSKEALLASQKPFKFIAREEQKRAAREKQLRDFLKYKKKTNRFKAR 358
Mm1   LVKQSEERRKKARERNRAALLASLKPFKFIAREEQKQAVREKKLRDLFRAKRKTNQFKAK 352
Mm2   LVKQSEERRKKARERNRAALLASLKPFKFIAREEQKQAVREKKLRDLFRAKRKTNQFKAK 352
Mm3   LVKQSEERRKKARERNRAALLASLKPFKFIAREEQKQAVREKKLRDLFRAKRKTNQFKAK 352
      ****.****:. :*:.: ***** ************:*.***:***::: *:***:***:

Hs1   PIPRSTYGSTTNDKLKEEELYRNLRTQLRAQEHLQNSSPLPCRSACGCRNPRCPEQAVKL 418
Hs2   PIPRSTYGSTTNDKLKEEELYRNLRTQLRAQEHLQNSSPLPCRSACGCRNPRCPEQAVKL 418
Mm1   PVPRFIYRPAASDKPKEEELYGDSRMLPKVRDLL-QNSPWPSRSACR--RFRDPRSPAKP 409
Mm2   PVPRFIYRPAASDKPKEEELYGDSRMLPKVRDLL-QNSPWPSRSACR--RFRDPRSPAKP 409
Mm3   PVPRFIYRPAASDKPKEEELYGDSRMLPKVRDLL-QNSPWPSRSACR--RFRDPRSPAKP 409
      *:**  *  ::.** ****** : *   :.:: * :.** *.****   . * *.. .*

Hs1   KCKHKVRCPTPDFEDLPERYQKHLSEHKSPKLLTVCKPFDLHASPHASIKREKILADIEA 478
Hs2   KCKHKVRCPTPDFEDLPERYQKHLSEHKSPKLLTVCKPFDLHASPHASIKREKILADIEA 478   UPF0564
Mm1   RGKHRRRCLRRDG--DLEKWKEPFSEYSFLKCPMLCEECCLHESPCDSDKRQKLLADIRA 467
Mm2   RGKHRRRCLRRDG--DLEKWKEPFSEYSFLKCPMLCEECCLHESPCDSDKRQKLLADIRA 467
Mm3   RGKHRRRCLRRDG--DLEKWKEPFSEYSFLKCPMLCEECCLHESPCDSDKRQKLLADIRA 467
      : **: **   *     *:::: :**:.  *   :*:   ** **  * **:*:****.*

Hs1   DEENLKETRWPYLSPRRKSPVRCAGVNPVPCNCNPPVPTVSSRGREQAVRRSLEEKKMLE 538
Hs2   DEENLKETRWPYLSPRRKSPVRCAGVNPVPCNCNPPVPTVSSRGREQAVR---------- 528
Mm1   DEEILRETRRPGRSPRRKSPGRSSNPKPRPHECSPPMPTASSRGREQAIRKSLEEKKMLE 527   Exon 4
Mm2   DEEILRETRRPGRSPRRKSPGRSSNPKPRPHECSPPMPTASSRGREQAIR---------- 517
Mm3   DEEILRETRRPGRSPRRKSPGRSSNPKPRPHECSPPMPTASSRGREQAIR---------- 517
      *** *:*** *  ******* *.:. :* * :*.**:**.*********:*

Hs1   EERNRILTKQKQRMKELQKLLTTRAKAYDSHQSLAQISKSRVKCLRKSEKERMREYQREL 598
Hs2   ----------------------------------------------KSEKERMREYQREL 542
Mm1   EERNRILTKQKQRMKDLQKLLATRVKAYGSHQSLSQIVKARVKELRRSEKARMREYWQEL 587
Mm2   ----------------------------------------------RSEKARMREYWQEL 531
Mm3   ----------------------------------------------RSEKARMREYWQEL 531
                                                    :*** ***** :**

Hs1   EEREEKLKKRPLLFERVAQKNARMAAEKHYSNTLKALGISDEFVSKKGQSGKVLEYFNNQ 658
Hs2   EEREEKLKKRPLLFERVAQKNARMAAEKHYSNTLKALGISDEFVSKKGQSGKVLEYFNNQ 602
Mm1   EEQEEKLQKRPMLFERVTQRNARMAAERCYSNTLKALGLLEEFVSEKGQSGKVSEDFTRQ 647
Mm2   EEQEEKLQKRPMLFERVTQRNARMAAERCYSNTLKALGLLEEFVSEKGQSGKVSEDFTRQ 591
Mm3   EEQEEKLQKRPMLFERVTQVVFIG------------------------------------ 555   Type 3 unique sequence
      **:****:***:******:*

Hs1   ETKSVTEDKESFNEEEKIEERENGEENYFIDTNSQDSYKEKDEANEES--EEEKSVEESH 716
Hs2   ETKSVTEDKESFNEEEKIEERENGEENYFIDTNSQDSYKEKDEANEES--EEEKSVEESH 660
Mm1   ELRSCTLEKESSYG----EERENEEESYLTDISSQDSCKGNKEDIKDIKSGEENSGE--- 700
Mm2   ELRSCTLEKESSYG----EERENEEESYLTDISSQDSCKGNKEDIKDIKSGEENSGE--- 644
Mm3   ------------------------------------------------------------ 555
```

**SFig 4-6 – POC5 localizes to canonical centrioles when each of the three FAM161A isoforms are overexpressed.**

**A**) POC5 localization relative to overexpressed human FAM161A (hFAM161A) type 2, house mouse FAM161A (mFAM161A) type 2, and mFAM161A type 3 in U2OS cells. Human FAM161A type 2 and mFAM161A type 2 exhibit centrosomal and pericentrosomal localization. **B**) Quantification showing the percentage of centrioles exhibiting centrosomal, pericentrosomal, and non-centrosomal localization of hFAM161A type 2, mFAM161A type 2, and mFAM161A type 3. **C**) POC5 localization relative to astral microtubules when overexpressing hFAM161A type 2, mFAM161A type 2, and mFAM161A type 3 in U2OS cells. **D**) Quantification showing the percentage of cells exhibiting centrosomal localization of POC5 based on astral microtubules. All experiments were repeated three times with consistent results.

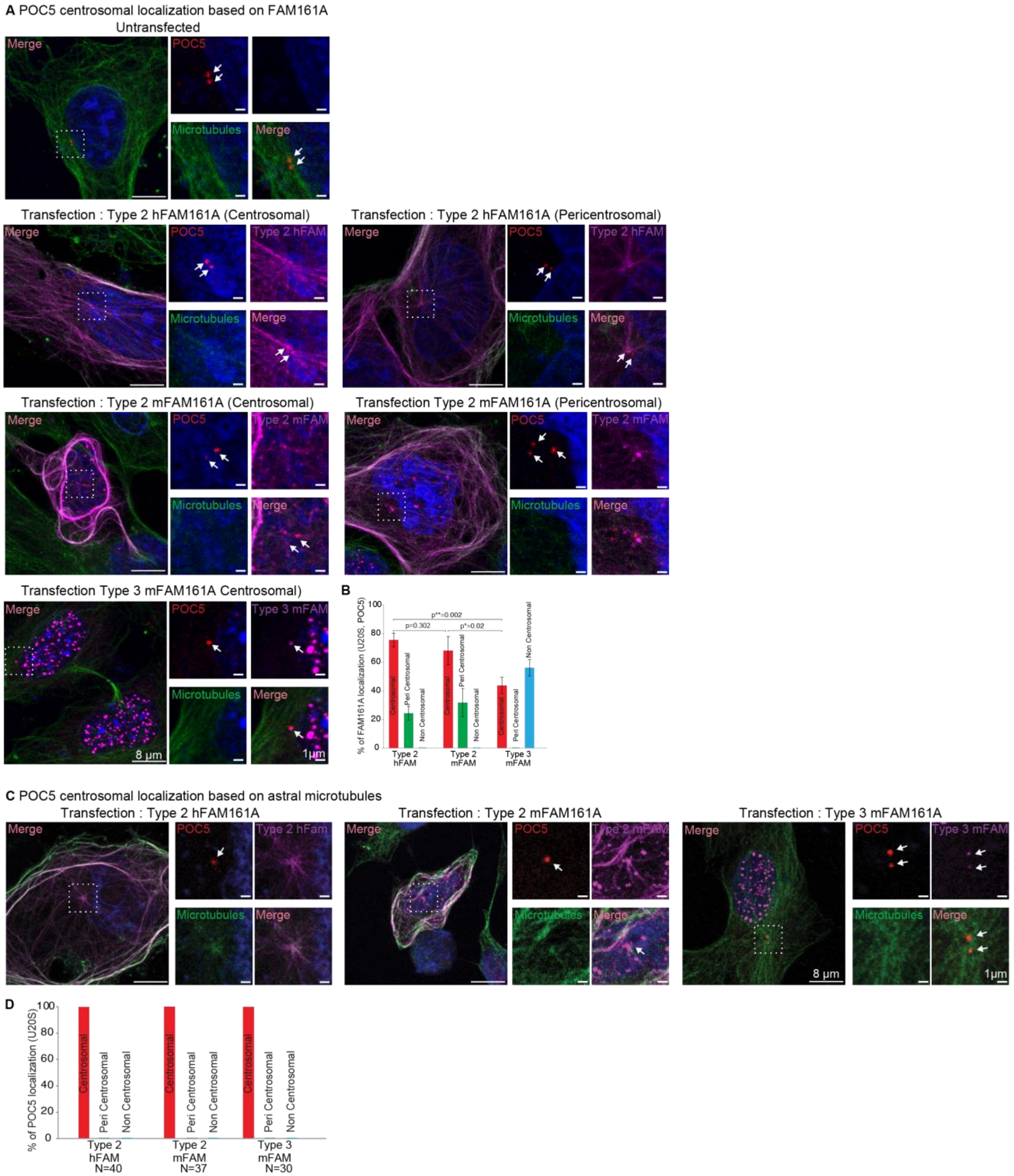
A POC5 centrosomal localization based on FAM161A
Untransfected
Merge
POC5
Microtubules
Merge
Transfection : Type 2 hFAM161A (Centrosomal)
Transfection : Type 2 hFAM161A (Pericentrosomal)
Type 2 hFAM
Transfection : Type 2 mFAM161A (Centrosomal)
Transfection Type 2 mFAM161A (Pericentrosomal)
Type 2 mFAM
Transfection Type 3 mFAM161A Centrosomal)
Type 3 mFAM
8 µm
1µm
B
% of FAM161A localization (U20S, POC5)
p**=0.002
p=0.302
p*=0.02
Centrosomal
Peri Centrosomal
Non Centrosomal
Type 2 hFAM
Type 2 mFAM
Type 3 mFAM
C POC5 centrosomal localization based on astral microtubules
Transfection : Type 2 hFAM161A
Transfection : Type 2 mFAM161A
Transfection : Type 3 mFAM161A
Type 2 hFam
D
% of POC5 localization (U20S)
Type 2 hFAM N=40
Type 2 mFAM N=37
Type 3 mFAM N=30

**SFig 4-7 – House mouse FAM161A type 3 inhibits type 2 from recruiting POC1B to the microtubules.**

**A–C**) Expression of human POC1B (hPOC1B) with house mouse FAM161A (mFAM161A) type 2 (**A**), mFAM161A type 3 (**B**), and both mFAM161A types 2 & 3 (**C**). Only one FAM161A isoform at a time is shown in panel C; however, the cells were transfected with all three proteins (i.e., hPOC1B and mFAM161A types 2 & 3). Scale bars are 8 µm. **D**) Quantification of hPOC1B colocalization with tubulin under various transfection conditions. **E**) mFAM161A type 3 and hPOC1B colocalize in extranuclear foci. Shown in inset. ****P<0.0001. n, number of cells. Scale bars are 5 µm. All images are representative of three independent experiments. The quantification data was compiled from three independent experiments.

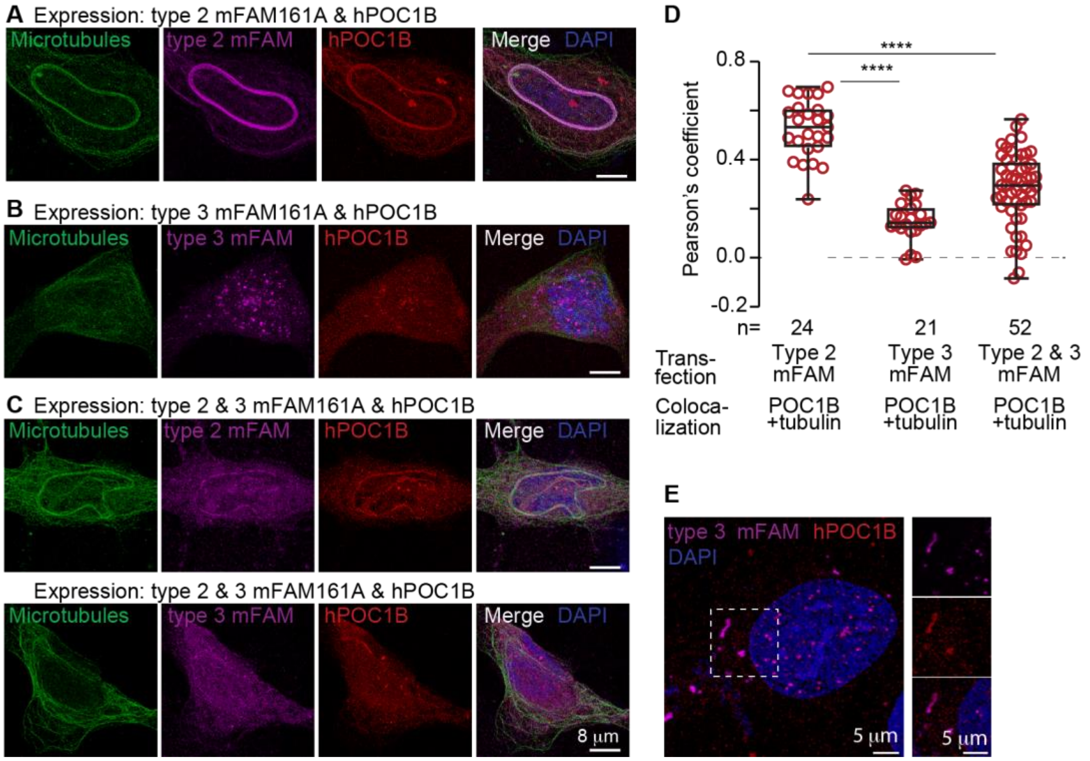

**SFig 5 – FAM161A labels the proximal and distal centrioles of dog spermatozoa.**

Confocal (deconvolution) microscopy of the dog distal centriole showed that FAM161A exhibits prominent left- and right-side labeling, suggesting that the distal centriolar rods are far away from each other. PC, proximal centriole; DC, distal centriole; Ax, axoneme. Scale bars are 1 µm. All images are representative of at least two independent experiments.

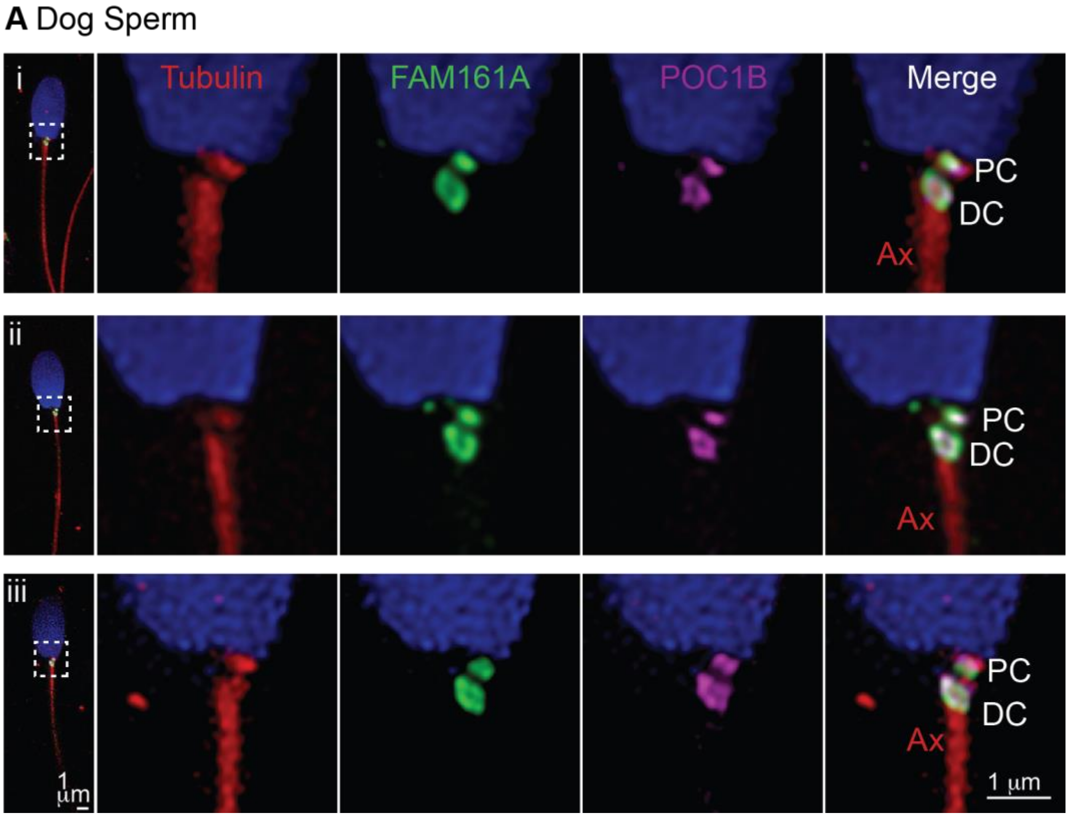

**SFig 6-1 – The bovine distal centriole begins splaying in round spermatids and splits in elongated spermatids.**

**A**) 3D-STORM imaging of bovine testes with rod protein staining during spermatogenesis. The white arrow marks the splayed centriole. **B**) Centriolar length in spermatogonia/spermatocytes (Sg), round spermatids (RS), and elongated spermatids (ES). The C1/2 panel includes both C1 and C2. **C**) Proximal and distal centriolar width at their caudal (aka tip) and rostral (aka base) ends in round spermatids. C1/2, centrioles 1 and 2; PC, proximal centriole; DC, distal centriole. **D**) Distal centriolar rod length and width measurements based on POC5 staining in elongated spermatids. All data shown were accumulated over three independent experiments. Data B and C are plotted as box and whisker with interquartile range. Data shown in **D** is an average ± SD. ****P<0.0001, ***P<0.001, **P<0.01, *P<0.05; ns, not significant; n, number of centrioles.

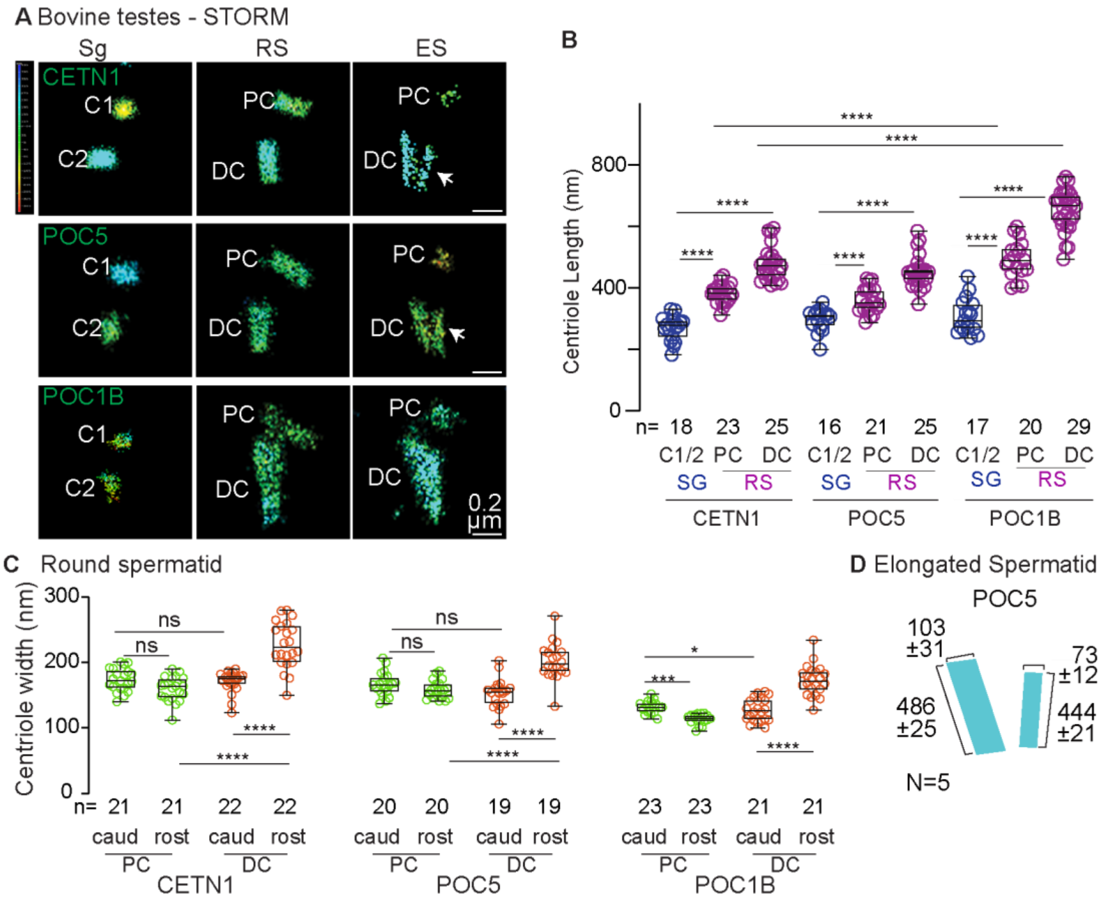

**SFig 6-2 – Centriole remodeling generates species-specific centriolar sizes.**

Quantification of centriolar length based on CETN1 labeling in bovine (Bt), rabbit (Oc), and human (Hs) testes at spermatogonia/spermatocyte (Sg) and round spermatid (RS) cell stages. C1/2, centriole 1/2; PC, proximal centriole; DC, distal centriole. ****P<0.0001, **P<0.01; ns, not significant; n, number of centrioles. The result is based on immunostaining with CETN1 and confocal imaging. The data was accumulated over three independent experiments.

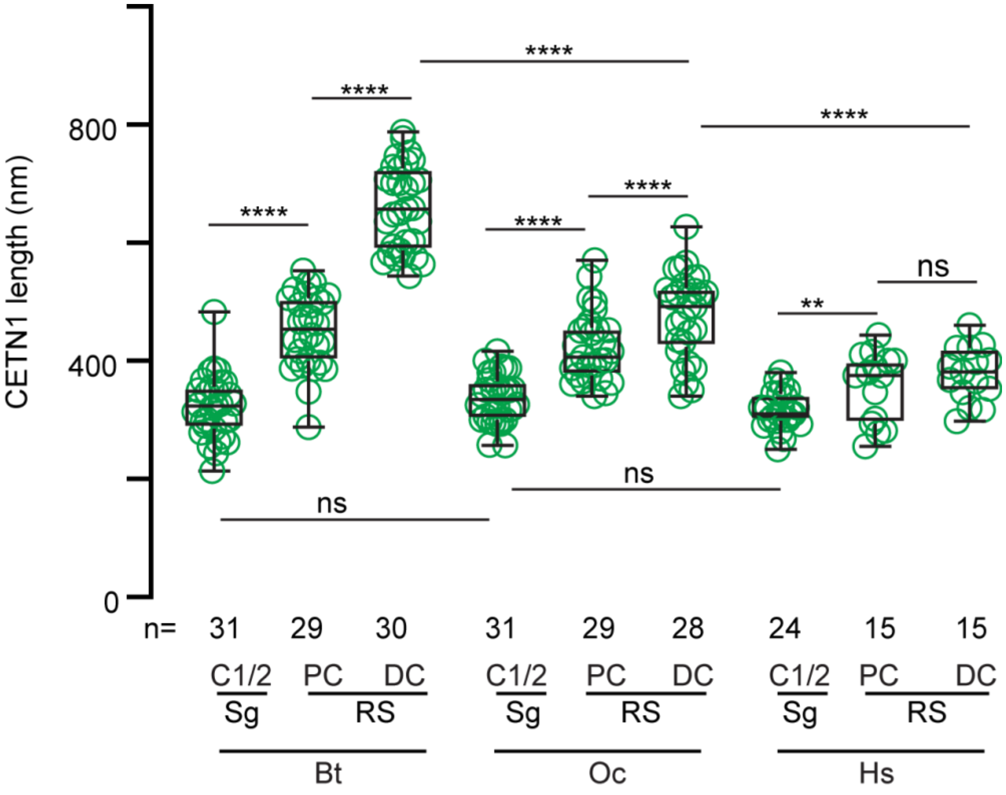

**SFig 6-3 – Quantification in the proximal and distal centrioles shows localization intensity changes similar to that observed during total centriole quantification.**

Quantification of various proteins in individual centrioles (C1 and C2) of spermatogonia/spermatocytes (Sg), and in proximal and distal centrioles (PC and DC) of round (RS) and elongated spermatids (ES) in bovine.. The graphs are shown as box and whisker plots with an interquartile range. ****P<0.0001, ***P<0.001, **P<0.01, *P<0.05; ns, not significant; n, number of cells.

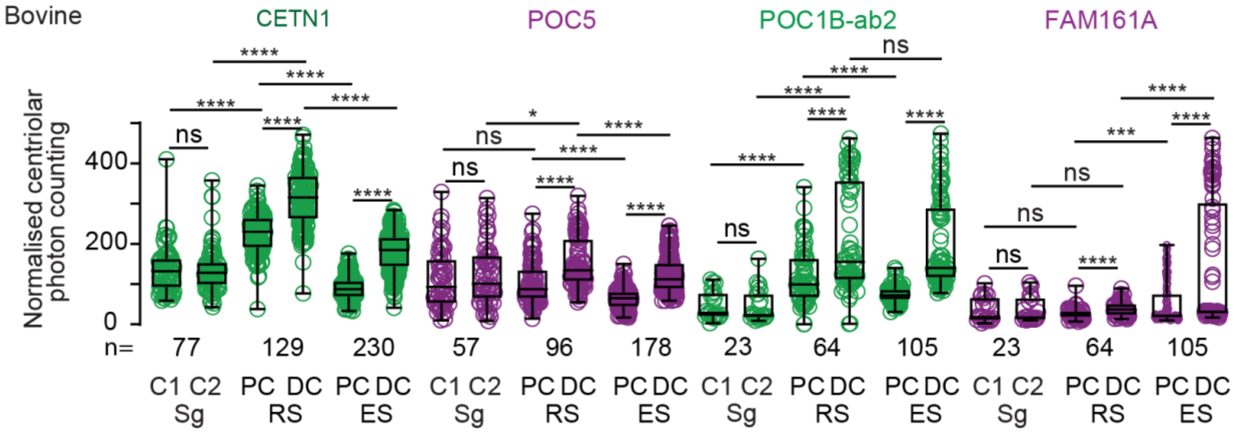

**SFig 7-1 – FAM161A N-terminus Ab labeling in mouse testes**

Immunostaining using FAM161A N-terminus antibody and POC1B ab2 in house mouse round spermatids (RS). PC, proximal centriole; DC, distal centriole. Scale bars are 1 µm. The result was consistent in three independent immunostainings.

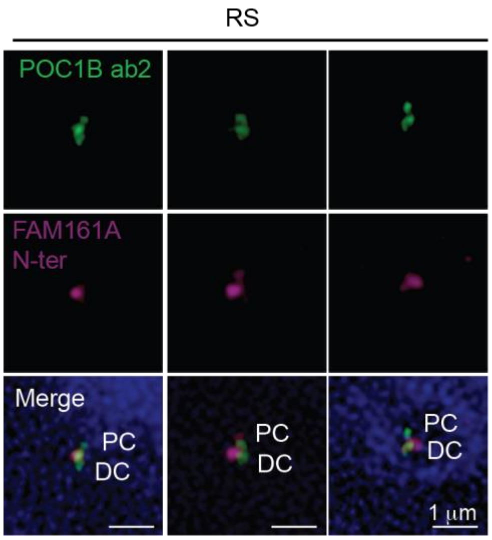

**SFig 7-2 – Quantification in the proximal and distal centrioles shows localization intensity changes similar to that observed during total centriole quantification.**

Quantification of various proteins in individual centrioles (C1 and C2) of spermatogonia/spermatocytes (Sg), and in proximal and distal centrioles (PC and DC) of round (RS) and elongated spermatids (ES) in mice. The distal centriole in the round spermatid was determined as the larger and brighter POC5-labeled centriole. The graphs are shown as box and whisker plots with an interquartile range. ****P<0.0001, ***P<0.001, **P<0.01, *P<0.05; ns, not significant; n, number of cells.

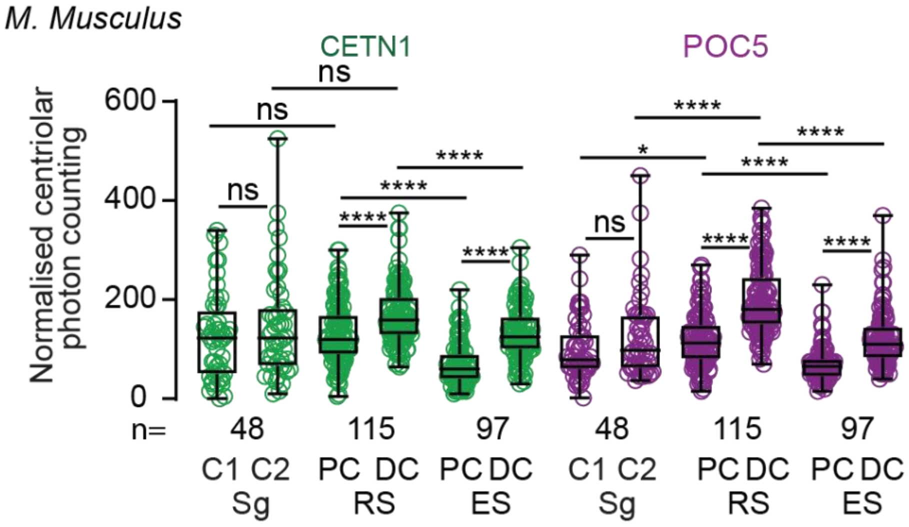

**STable 8-1 – Antibodies used in this study**

| Antibody<br>(Production animal) | Company<br>(Catalog number) | Concentration (Application) |
|---|---|---|
| POC5 ab1 (Rabbit) | Gifted by Dr. Bernan (PMID #19349582) | 1:200 (Confocal)<br>1:100 (STORM) |
| POC5 ab2 (Rabbit) | Thermo Fisher Scientific (PA5-24308) | 1:200 (Confocal)<br>1:100 (STORM) |
| CETN1 (Mouse) | Santa Cruz (2A6) | 1:10 (Confocal)<br>1:5 (STORM) |
| POC1B ab1 (Rabbit) | Thermo Fisher Scientific (PA5-24495) | 1:300 (Confocal) |
| POC1B ab2 (Mouse) | Thermo Fisher Scientific (H00282809-B01P) | 1:300 (Confocal)<br>1:100 (STORM) |
| FAM161A ab1 (Rabbit) | Sigma Aldrich (HPA032119) | 1:300 (Confocal) |
| FAM161A ab2 (Rabbit) | Novus Biologicals (NBP1-91508) | 1:1000 (Western) |
| FAM161A N-terminus (Rabbit) | Gifted by Dr. Dror Sharon (Hadassah-Hebrew University Medical Center) | 1:200 (Confocal) |
| Tubulin (Mouse) | DSHB (Developmental Studies Hybridoma Bank) (E7) | 1:20 (Supernatant)<br>1:100 (Concentrate) |
| Tubulin (Sheep) | Cytoskeleton, Inc. (ATN02) | 1:500 (Confocal) |
| Anti-HA (Rabbit) | Invitrogen (SG77) | 1:200 (Confocal)<br>1:1000 (Western) |
| Anti-FLAG (Mouse) | Invitrogen (FG4R) | 1:400 (Confocal)<br>1:1000 (Western) |
| Anti-Rabbit A647 (Donkey) | Jackson ImmunoResearch (711-605-152) | 1:300 (Confocal)<br>1:100 (STORM) |
| Anti-Mouse A647 (Donkey) | Jackson ImmunoResearch (715-605-150) | 1:300 (Confocal)<br>1:100 (STORM) |
| Anti-Rabbit A488 (Donkey) | Jackson ImmunoResearch (711-545-152) | 1:200 (Confocal) |
| Anti-Sheep A555 (Donkey) | Thermo Fisher Scientific (A-21436) | 1:500 (Confocal) |
| Anti-Mouse A488 (Donkey) | Jackson ImmunoResearch (715-545-150) | 1:200 (Confocal) |
| Anti-Sheep A488 (Donkey) | Jackson ImmunoResearch (715-545-003) | 1:300 (Confocal) |

**STable 8-2 – Primers used in this study**

| Application | Primer | Sequence |
|---|---|---|
| 3’ RACE | Oligo d(T)-Anchor | 5’-GACCACGCGTATCGATGTCGACTTTTTTTTTTTTTTTTV-3’<br>*Mlu* **I** site *Cla* **I** site *Sal* **I** site, V = A, C, or G (what is that) |
| 1$^{st}$ PCR | 2FW | 5’-CAGGAAACTCAAAGACCTGAAGG-3’ |
| | PCR Anchor | 5’-GACCACGCGTATCGATGTCGAC-3’ |
| 2$^{nd}$ PCR | 3FW | 5’-GCCAGCTGCGAGTGACAAGC-3’ |
| | 5RV | 5’-CTGGAGGAACAAGAGGAAAAACTGC-3’ |
| | 5*RV | 5’-GCCTCCACCTTTGTGCTCCGTC-3’ |
| | 7RV | 5’-GAACCACAGCTGACACAAATGG-3’ |
| Sequencing | Seq. 3FW | 5’-GCAGACATCAGAGCAGATGAAG-3’ |
| | Seq. 5RV | 5’-GGAGTGAAAAGGCCAGGATGAG-3’ |
| | Seq. 5*RV | 5’-CCAGCTTTCTGTTTCTCAGTAC-3’ |
| | Seq. 7RV | 5’-GGAGAAGAAAGAGAGAATGAGG-3’ |

**STable 8-3 – Species analyzed in this study**

Table 8-3a. Rodent sequences analyzed in this study

| Organism | Length of amino acid sequence | Accession ID |
|---|---|---|
| **Mus musculus**<br>house mouse | **700** | XM_006514828.5 |
| **Rattus norvegicus**<br>Norway rat | **653** | XM_017599131.2 |
| **Heterocephalus glaber**<br>naked mole-rat | **706** | XM_004867228.3 |
| **Nannospalax galili**<br>Upper Galilee Mountains blind mole rat | **709** | XM_017800967.2 |
| **Mesocricetus auratus**<br>golden hamster | **695** | XM_005070431.4 |
| **Dipodomys ordii**<br>Ord’s kangaroo rat | **713** | XM_013022573.1 |
| **Castor canadensis**<br>American beaver | **701** | XM_020175016.1 |
| **Mus caroli**<br>Ryukyu mouse | **728** | XM_029483590.1 |
| **Mus pahari**<br>shrew mouse | **550** | XM_029545237.1 |
| **Peromyscus maniculatus bairdii**<br>prairie deer mouse | **699** | XM_006974797.2 |
| **Jaculus jaculus**<br>lesser Egyptian jerboa | **921** | XM_004665507.1 |
| **Meriones unguiculatus**<br>Mongolian gerbil | **699** | XM_021626659.1 |
| **Rattus rattus**<br>black rat | **471** | XM_032916924.1 |
| **Grammomys surdaster**<br>thicket rat | **260** | XM_028778791.1 |
| **Mastomys coucha**<br>southern multimammate mouse | **495** | XM_031341979.1 |
| **Arvicanthis niloticus**<br>African grass rat | **641** | XM_034508367.1 |
| **Cricetulus griseus**<br>Chinese hamster | **703** | XM_003502153.5 |
| **Peromyscus leucopus**<br>white-footed mouse | **782** | XM_028876357.2 |
| **Onychomys torridus**<br>southern grasshopper mouse | **696** | XM_036201671.1 |
| **Arvicola amphibius**<br>Eurasian water vole | **704** | XM_038339639.1 |
| **Microtus oregoni**<br>creeping vole | **705** | XM_041669288.1 |
| **Microtus ochrogaster**<br>prairie vole | **706** | XM_026788902.1 |
| **Cavia porcellus**<br>domestic guinea pig | **686** | XM_013150060.2 |
| **Octodon degus**<br>degu | **702** | XM_023720875.1 |
| **Fukomys damarensis**<br>Damaraland mole-rat | **693** | XM_033762286.1 |
| **Marmota marmota**<br>alpine marmot | **551** | XM_015479628.1 |
| **Ictidomys tridecemlineatus**<br>thirteen-lined ground squirrel | **705** | XM_005322047.4 |
| **Urocitellus parryii** | **705** | XM_026387235.1 |

| | | |
|---|---|---|
| Arctic ground squirrel | | |
| **Marmota flaviventris**<br>yellow-bellied marmot | **705** | XM_027934553.1 |

Table 8-3b. Carnivore sequences analyzed in this study.

| Organism | Length of amino acid sequence | Accession ID |
|---|---|---|
| **Canis lupus familiaris**<br>dog | 716 | XM_005626140.4 |
| **Mustela putorius furo**<br>domestic ferret | 729 | XM_004742028.2 |
| **Panthera tigris altaica**<br>Amur tiger | 665 | XM_007075417.1 |
| **Enhydra lutris kenyoni**<br>northern sea otter | 720 | XM_022504396.1 |
| **Acinonyx jubatus**<br>cheetah | 715 | XM_027072376.1 |
| **Lynx canadensis**<br>Canada lynx | 715 | XM_030310672.1 |
| **Puma yagouaroundi**<br>jaguarundi | 715 | XM_040463327.1 |
| **Puma concolor**<br>puma | 714 | XM_025932404.1 |
| **Felis catus**<br>domestic cat | 715 | XM_003984033.5 |
| **Panthera pardus**<br>leopard | 715 | XM_019431779.1 |
| **Canis lupus dingo**<br>dingo | 716 | XM_025468782.2 |
| **Ursus arctos horribilis**<br>grizzly bear | 719 | XM_026485066.1 |
| **Lontra canadensis**<br>North American river otter | 721 | XM_032873831.1 |
| **Mustela erminea**<br>ermine | 729 | XM_032353647.1 |
| **Odobenus rosmarus divergens**<br>Pacific walrus | 719 | XM_004397721.1 |
| **Hyaena hyaena**<br>striped hyena | 705 | XM_039240346.1 |
| **Suricata suricatta**<br>meerkat | 706 | XM_029937442.1 |
| **Vulpes lagopus**<br>Arctic fox | | XM_041757534.1 |
| **Vulpes vulpes**<br>red fox | 776 | XM_025992735.1 |
| **Ursus maritimus**<br>polar bear | 719 | XM_040623440.1 |
| **Ailuropoda melanoleuca**<br>giant panda | 720 | XM_034659197.1 |
| **Eumetopias jubatus**<br>Steller sea lion | 718 | XM_028089669.1 |
| **Zalophus californianus**<br>California sea lion | 718 | XM_027624320.1 |
| **Neomonachus schauinslandi**<br>Hawaiian monk seal | 721 | XM_021699187.1 |
| **Halichoerus grypus**<br>grey seal | 722 | XM_036121192.1 |

| | | |
|---|---|---|
| **Leptonychotes weddellii**<br>Weddell seal | 722 | XM_006733195.2 |
| **Mirounga leonina**<br>southern elephant seal | 720 | XM_035001465.1 |
| **Phoca vitulina**<br>harbor seal | 722 | XM_032410896.1 |

Table 8-3c. Ungulate sequences analyzed in this study.

| Organism | Length of amino acid sequence | Accession ID |
|---|---|---|
| **Bos taurus**<br>cattle | 721 | XM_005212825.4 |
| **Odocoileus virginianus texanus**<br>Texas whitetail deer | 716 | XM_020891631.1 |
| **Bison bison**<br>American bison | 721 | XM_010863278.1 |
| **Bos indicus x Bos taurus**<br>hybrid cattle | 721 | XM_027555592.1 |
| **Bos mutus**<br>wild yak | 717 | XM_005889728.2 |
| **Bos indicus**<br>zebu | 710 | XM_019969488.1 |
| **Bubalus bubalis**<br>water buffalo | 721 | XM_006049386.2 |
| **Oryx dammah**<br>scimitar-horned oryx | 810 | XM_040224006.1 |
| **Capra hircus**<br>goat | 722 | XM_013967738.2 |
| **Ovis aries**<br>sheep | 722 | XM_027966713.2 |
| **Sus scrofa**<br>pig | 720 | XM_003354802.4 |
| **Vicugna pacos**<br>alpaca | 723 | XM_015237028.2 |
| **Camelus ferus**<br>wild Bactrian camel | 723 | XM_014564859.2 |
| **Camelus bactrianus**<br>Bactrian camel | 552 | XM_010951902.1 |
| **Camelus dromedarius**<br>Arabian camel | 722 | XM_010989773.2 |
| **Neophocaena asiaeorientalis**<br>Yangtze finless porpoise | 713 | XM_024753025.1 |
| **Balaenoptera acutorostrata scammony**<br>minke whale | 717 | XM_007189708.1 |
| **Lipotes vexillifer**<br>Yangtze River dolphin | 713 | XM_007464134.1 |
| **Lagenorhynchus obliquidens**<br>Pacific white-sided dolphin | 717 | XM_027131791.1 |
| **Globicephala melas**<br>long-finned pilot whale | 717 | XM_030877335.1 |
| **Orcinus orca**<br>killer whale | 717 | XM_004280649.2 |
| **Tursiops truncatus**<br>common bottlenose dolphin | 717 | XM_019943077.2 |
| **Phocoena sinus**<br>vaquita | 713 | XM_032653378.1 |

| | | |
|---|---|---|
| **Monodon monoceros**<br>narwhal | 717 | XM_029232453.1 |
| **Delphinapterus leucas**<br>beluga whale | 717 | XM_022597296.2 |
| **Physeter catodon**<br>sperm whale | 727 | XM_024133601.2 |
| **Balaenoptera musculus**<br>blue whale | **727** | XM_036873937.1 |

Table 8-3d. Primate sequences analyzed in this study.

| Organism | Length of amino acid sequence | Accession ID |
|---|---|---|
| **Homo sapiens**<br>human | 716 | NM_001201543.2 |
| **Pan troglodytes**<br>chimpanzee | 716 | XM_515502.6 |
| **Macaca mulatta**<br>rhesus macaque | 716 | XM_028831643.1 |
| **Macaca fascicularis**<br>crab-eating macaque | 716 | XM_005575777.2 |
| **Callithrix jacchus**<br>white-tufted-ear marmoset | 715 | XM_003734729.4 |
| **Colobus angolensis palliates**<br>Angola black-and-white colobus | 562 | XM_011941831.1 |
| **Chlorocebus sabaeus**<br>green monkey | 716 | XM_007970494.2 |
| **Cercocebus atys**<br>sooty mangabey | 714 | XM_012039548.1 |
| **Macaca nemestrina**<br>pig-tailed macaque | 716 | XM_011713355.2 |
| **Papio anubis**<br>olive baboon | 716 | XM_003908708.5 |
| **Theropithecus gelada**<br>gelada | 716 | XM_025354341.1 |
| **Mandrillus leucophaeus**<br>drill | 716 | XM_011987312.1 |
| **Trachypithecus francoisi**<br>François' langur | 711 | XM_033239161.1 |
| **Rhinopithecus bieti**<br>black snub-nosed monkey | 711 | XM_017889712.1 |
| **Rhinopithecus roxellana**<br>golden snub-nosed monkey | 711 | XM_010384442.2 |
| **Piliocolobus tephrosceles**<br>Ugandan red colobus | 711 | XM_023228337.1 |
| **Gorilla gorilla**<br>western gorilla | 716 | XM_004029293.3 |
| **Pan paniscus**<br>pygmy chimpanzee | 716 | XM_003830874.3 |
| **Pongo abelii**<br>Sumatran orangutan | 716 | NM_001374141.1 |
| **Nomascus leucogenys**<br>northern white-cheeked gibbon | 716 | XM_003262434.4 |
| **Hylobates moloch**<br>silvery gibbon | 716 | XM_032177370.1 |
| **Saimiri boliviensis**<br>Bolivian squirrel monkey | 715 | XM_039464018.1 |
| **Sapajus apella**<br>tufted capuchin | 715 | XM_032292015.1 |
| **Cebus imitator**<br>Panamanian white-faced capuchin | 715 | XM_017522196.2 |
| **Aotus nancymaae**<br>Ma’s night monkey | 715 | XM_012477168.2 |
| **Carlito syrichta**<br>Philippine tarsier | 715 | XM_008054794.2 |
| **Propithecus coquereli**<br>Coquerel’s sifaka | 714 | XM_012639195.1 |
| **Microcebus murinus** | 708 | XM_012764864.2 |

| | | |
|---|---|---|
| gray mouse lemur | | |
| **Otolemur garnettii**<br>small-eared galago | 712 | XM_003787945.1 |

Le Guennec, M., N. Klena, D. Gambarotto, M.H. Laporte, A.M. Tassin, H. van den Hoek, P.S. Erdmann, M. Schaffer, L. Kovacik, S. Borgers, K.N. Goldie, H. Stahlberg, M. Bornens, J. Azimzadeh, B.D. Engel, V. Hamel, and P. Guichard. 2020. A helical inner scaffold provides a structural basis for centriole cohesion. *Sci Adv*. 6:eaaz4137.

Lee, J.-H., and T. Mori. 2006. Ultrastructural observations on the sperm of two Apodemus species, Apodemus agrarius coreae and Apodemus speciosus peninsulae, in Korea.

Lee, J.-H., and K.-R. Park. 2011. Fine Structure of Sperm in the Korea Squirrel, Tamias sibiricus. *Applied Microscopy*. 41:99-107.

Lemaitre, J.F., J.M. Gaillard, and S.A. Ramm. 2020. The hidden ageing costs of sperm competition. *Ecol Lett*. 23:1573-1588.

Leung, M.R., M.C. Roelofs, R.T. Ravi, P. Maitan, H. Henning, M. Zhang, E.G. Bromfield, S.C. Howes, B.M. Gadella, H. Bloomfield-Gadelha, and T. Zeev-Ben-Mordehai. 2021. The multi-scale architecture of mammalian sperm flagella and implications for ciliary motility. *EMBO J*. 40:e107410.

Li, Y.Z., N. Li, W.S. Liu, Y.W. Sha, R.F. Wu, Y.L. Tang, X.S. Zhu, X.L. Wei, X.Y. Zhang, Y.F. Wang, Z.X. Lu, and F.X. Zhang. 2022. Biallelic mutations in spermatogenesis and centriole-associated 1 like

(SPATC1L) cause acephalic spermatozoa syndrome and male infertility. *Asian journal of andrology*. 24:67-72.

Luke, L., M. Tourmente, and E.R. Roldan. 2016. Sexual Selection of Protamine 1 in Mammals. *Mol Biol Evol*. 33:174-184.

Lüpold, S., R.A. de Boer, J.P. Evans, J.L. Tomkins, and J.L. Fitzpatrick. 2020. How sperm competition shapes the evolution of testes and sperm: a meta-analysis. *Philosophical Transactions of the Royal Society B*. 375:20200064.

Lüpold, S., and S. Pitnick. 2018. Sperm form and function: what do we know about the role of sexual selection? *Reproduction*. 155:R229-R243.

Manandhar, G., H. Schatten, and P. Sutovsky. 2005. Centrosome reduction during gametogenesis and its significance. *Biology of reproduction*. 72:2-13.

Manandhar, G., C. Simerly, J.L. Salisbury, and G. Schatten. 1999. Centriole and centrin degeneration during mouse spermiogenesis. *Cell Motil Cytoskeleton*. 43:137-144.

Manandhar, G., C. Simerly, and G. Schatten. 2000. Highly degenerated distal centrioles in rhesus and human spermatozoa. *Hum Reprod*. 15:256-263.

Manandhar, G., P. Sutovsky, H.C. Joshi, T. Stearns, and G. Schatten. 1998. Centrosome reduction during mouse spermiogenesis. *Dev Biol*. 203:424-434.

Mercey, O., C. Kostic, E. Bertiaux, A. Giroud, Y. Sadian, D.C. Gaboriau, C.G. Morrison, N. Chang, Y. Arsenijevic, and P. Guichard. 2022. The connecting cilium inner scaffold provides a structural foundation that protects against retinal degeneration. *PLoS Biology*. 20:e3001649.

Mokos, J., I. Scheuring, A. Liker, R.P. Freckleton, and T. Székely. 2021. Degree of anisogamy is unrelated to the intensity of sexual selection. *Scientific reports*. 11:1-11.

Murrell, B., J.O. Wertheim, S. Moola, T. Weighill, K. Scheffler, and S.L. Kosakovsky Pond. 2012. Detecting individual sites subject to episodic diversifying selection. *PLoS Genet*. 8:e1002764.

Palacios Martinez, S., J. Greaney, and J. Zenker. 2022. Beyond the centrosome: The mystery of microtubule organising centres across mammalian preimplantation embryos. *Curr Opin Cell Biol*. 77:102114.

Parker, G.A. 1970. Sperm Competition and Its Evolutionary Consequences in the Insects. *Biological Reviews*. 45:525-567.

Parker, G.A. 2014. The sexual cascade and the rise of pre-ejaculatory (Darwinian) sexual selection, sex roles, and sexual conflict. *Cold Spring Harbor perspectives in biology*. 6:a017509.

Phillips, D.M. 1967. Giant centriole formation in Sciara. *The Journal of cell biology*. 33:73-92.

Plant, T.M., and A.J. Zeleznik. 2014. Knobil and Neill's physiology of reproduction. Academic Press.

Pomp, O., H.Y.G. Lim, R.M. Skory, A.A. Moverley, P. Tetlak, S. Bissiere, and N. Plachta. 2022. A monoastral mitotic spindle determines lineage fate and position in the mouse embryo. *Nature cell biology*. 24:155-167.

Rambaut, A. University of Edinburgh, Institute of Evolutionary Biology; Edinburgh, UK: 2012. FigTree V. 1.4. Molecular Evolution, Phylogenetics and Epidemiology.

Rawe, V.Y., Y. Terada, S. Nakamura, C.F. Chillik, S.B. Olmedo, and H.E. Chemes. 2002. A pathology of the sperm centriole responsible for defective sperm aster formation, syngamy and cleavage. *Hum Reprod*. 17:2344-2349.

Roldan, E.R., M. Gomendio, and A.D. Vitullo. 1992. The evolution of eutherian spermatozoa and underlying selective forces: female selection and sperm competition. *Biol Rev Camb Philos Soc*. 67:551-593.

Roldan, E.R.S. 2019. Sperm competition and the evolution of sperm form and function in mammals. *Reprod Domest Anim*. 54 Suppl 4:14-21.

Ronquist, F., M. Teslenko, P. van der Mark, D.L. Ayres, A. Darling, S. Hohna, B. Larget, L. Liu, M.A. Suchard, and J.P. Huelsenbeck. 2012. MrBayes 3.2: efficient Bayesian phylogenetic inference and model choice across a large model space. *Syst Biol*. 61:539-542.

Roosing, S., I.J. Lamers, E. de Vrieze, L.I. van den Born, S. Lambertus, H.H. Arts, P.B.S. Group, T.A. Peters, C.B. Hoyng, H. Kremer, L. Hetterschijt, S.J. Letteboer, E. van Wijk, R. Roepman, A.I. den Hollander, and F.P. Cremers. 2014. Disruption of the basal body protein POC1B results in autosomal-recessive cone-rod dystrophy. *Am J Hum Genet*. 95:131-142.

Ross, B.D., L. Rosin, A.W. Thomae, M.A. Hiatt, D. Vermaak, A.F. de la Cruz, A. Imhof, B.G. Mellone, and H.S. Malik. 2013. Stepwise evolution of essential centromere function in a Drosophila neogene. *Science*. 340:1211-1214.

Rowley, A.G., T.S. Daly-Engel, and J.L. Fitzpatrick. 2019. Testes size increases with sperm competition risk and intensity in bony fish and sharks. *Behavioral Ecology*. 30:364-371.

Santos, P.R., M.F. Oliveira, M.A. Arroyo, A.R. Silva, R.E. Rici, M.A. Miglino, and A.C. Assis Neto. 2014. Ultrastructure of spermatogenesis in Spix's yellow-toothed cavy (Galea spixii). *Reproduction*. 147:13-19.

Schatten, G., C. Simerly, and H. Schatten. 1985. Microtubule configurations during fertilization, mitosis, and early development in the mouse and the requirement for egg microtubule-mediated motility during mammalian fertilization. *Proc Natl Acad Sci U S A*. 82:4152-4156.

Schatten, H., G. Schatten, D. Mazia, R. Balczon, and C. Simerly. 1986. Behavior of centrosomes during fertilization and cell division in mouse oocytes and in sea urchin eggs. *Proc Natl Acad Sci U S A*. 83:105-109.

Scheffler, K., J. Uraji, I. Jentoft, T. Cavazza, E. Monnich, B. Mogessie, and M. Schuh. 2021. Two mechanisms drive pronuclear migration in mouse zygotes. *Nat Commun*. 12:841.

Schill, D.J., G.W. LaBar, E.R.J.M. Mamer, and K.A. Meyer. 2010. Sex Ratio, Fecundity, and Models Predicting Length at Sexual Maturity of Redband Trout in Idaho Desert Streams. *North American Journal of Fisheries Management*. 30:1352-1363.

Shin, T.-Y., Y. Noguchi, Y. Yamamoto, K. MOCHIDA, and A. OGURA. 1998. Microtubule organization in hamster oocytes after fertilization with mature spermatozoa and round spermatids. *Journal of Reproduction and Development*. 44:185-189.

Simerly, C.R., N.B. Hecht, E. Goldberg, and G. Schatten. 1993. Tracing the incorporation of the sperm tail in the mouse zygote and early embryo using an anti-testicular alpha-tubulin antibody. *Dev Biol*. 158:536-548.

Soley, J.T. 2016. A comparative overview of the sperm centriolar complex in mammals and birds: Variations on a theme. *Animal reproduction science*. 169:14-23.

Steib, E., M.H. Laporte, D. Gambarotto, N. Olieric, C. Zheng, S. Borgers, V. Olieric, M. Le Guennec, F. Koll, A.M. Tassin, M.O. Steinmetz, P. Guichard, and V. Hamel. 2020. WDR90 is a centriolar microtubule wall protein important for centriole architecture integrity. *eLife*. 9:e57205.

Steppan, S., R. Adkins, and J. Anderson. 2004. Phylogeny and divergence-date estimates of rapid radiations in muroid rodents based on multiple nuclear genes. *Syst Biol*. 53:533-553.

Steppan, S.J., and J.J. Schenk. 2017. Muroid rodent phylogenetics: 900-species tree reveals increasing diversification rates. *PLoS One*. 12:e0183070.

Sutovsky, P., and G. Schatten. 2000. Paternal contributions to the mammalian zygote: fertilization after sperm-egg fusion. *Int Rev Cytol*. 195:1-65.

Swanson, W.J., and V.D. Vacquier. 2002. The rapid evolution of reproductive proteins. *Nature reviews. Genetics*. 3:137-144.

Tapia Contreras, C., and S. Hoyer-Fender. 2021. The Transformation of the Centrosome into the Basal Body: Similarities and Dissimilarities between Somatic and Male Germ Cells and Their Relevance for Male Fertility. *Cells*. 10.

Temple-Smith, P., A. Ravichandran, and F. Horta. 2018. Sperm: comparative vertebrate. *Encyclopedia of Reproduction*. 2:210-220.

Thybert, D., M. Roller, F.C.P. Navarro, I. Fiddes, I. Streeter, C. Feig, D. Martin-Galvez, M. Kolmogorov, V. Janousek, W. Akanni, B. Aken, S. Aldridge, V. Chakrapani, W. Chow, L. Clarke, C. Cummins, A. Doran, M. Dunn, L. Goodstadt, K. Howe, M. Howell, A.A. Josselin, R.C. Karn, C.M. Laukaitis, L. Jingtao, F. Martin, M. Muffato, S. Nachtweide, M.A. Quail, C. Sisu, M. Stanke, K. Stefflova, C. Van Oosterhout, F. Veyrunes, B. Ward, F. Yang, G. Yazdanifar, A. Zadissa, D.J. Adams, A. Brazma, M. Gerstein, B. Paten, S. Pham, T.M. Keane, D.T. Odom, and P. Flicek. 2018. Repeat associated mechanisms of genome evolution and function revealed by the Mus caroli and Mus pahari genomes. *Genome Res*. 28:448-459.

Toll-Riera, M., S. Laurie, and M.M. Alba. 2011. Lineage-specific variation in intensity of natural selection in mammals. *Mol Biol Evol*. 28:383-398.

Tung, C.K., and S.S. Suarez. 2021. Co-Adaptation of Physical Attributes of the Mammalian Female Reproductive Tract and Sperm to Facilitate Fertilization. *Cells*. 10:1297.

Turner, K., N. Solanki, H.O. Salouha, and T. Avidor-Reiss. 2022. Atypical Centriolar Composition Correlates with Internal Fertilization in Fish. *Cells*. 11:758.

Uzbekov, R., G. Singina, E. Shedova, C. Banliat, T. Avidor-Reiss, and S. Uzbekova. 2023. Centrosome formation in the bovine early embryo. *bioRxiv*:2022.2011.2029.517493.

Van Der Horst, G., L. Maree, S.H. Kotze, and M.J. O'Riain. 2011. Sperm structure and motility in the eusocial naked mole-rat, Heterocephalus glaber: a case of degenerative orthogenesis in the absence of sperm competition? *BMC Evol Biol*. 11:351.

Varea Sanchez, M., M. Bastir, and E.R. Roldan. 2013. Geometric morphometrics of rodent sperm head shape. *PLoS One*. 8:e80607.

Wang, G., Y. Guo, T. Zhou, X. Shi, J. Yu, Y. Yang, Y. Wu, J. Wang, M. Liu, X. Chen, W. Tu, Y. Zeng, M. Jiang, S. Li, P. Zhang, Q. Zhou, B. Zheng, C. Yu, Z. Zhou, X. Guo, and J. Sha. 2013. In-depth proteomic analysis of the human sperm reveals complex protein compositions. *Journal of proteomics*. 79:114-122.

Winey, M., and E. O'Toole. 2014. Centriole structure. *Philosophical transactions of the Royal Society of London. Series B, Biological sciences*. 369:20130457.

Woolley, D.M., and D.W. Fawcett. 1973. The degeneration and disappearance of the centrioles during the development of the rat spermatozoon. *Anat Rec*. 177:289-301.

Wu, B., H. Gao, C. Liu, and W. Li. 2020. The coupling apparatus of the sperm head and taildagger. *Biology of reproduction*. 102:988-998.

Xu, B., Z. Hao, K.N. Jha, Z. Zhang, C. Urekar, L. Digilio, S. Pulido, J.F. Strauss, 3rd, C.J. Flickinger, and J.C. Herr. 2008. TSKS concentrates in spermatid centrioles during flagellogenesis. *Dev Biol*. 319:201-210.

Yabe, T., X. Ge, and F. Pelegri. 2007. The zebrafish maternal-effect gene cellular atoll encodes the centriolar component sas-6 and defects in its paternal function promote whole genome duplication. *Dev Biol*. 312:44-60.

Yamauchi, Y., R. Yanagimachi, and T. Horiuchi. 2002. Full-term development of golden hamster oocytes following intracytoplasmic sperm head injection. *Biology of reproduction*. 67:534-539.

Yan, W., K. Morozumi, J. Zhang, S. Ro, C. Park, and R. Yanagimachi. 2008. Birth of mice after intracytoplasmic injection of single purified sperm nuclei and detection of messenger RNAs and MicroRNAs in the sperm nuclei. *Biology of reproduction*. 78:896-902.

Yanagimachi, R., Y. Kamiguchi, S. Sugawara, and K. Mikamo. 1983. Gametes and fertilization in the Chinese hamster. *Gamete research*. 8:97-117.

Yang, Z. 2007. PAML 4: phylogenetic analysis by maximum likelihood. *Mol Biol Evol*. 24:1586-1591.

Zach, F., F. Grassmann, T. Langmann, N. Sorusch, U. Wolfrum, and H. Stohr. 2012. The retinitis pigmentosa 28 protein FAM161A is a novel ciliary protein involved in intermolecular protein interaction and microtubule association. *Hum Mol Genet*. 21:4573-4586.

Zach, F., and H. Stohr. 2014. FAM161A, a novel centrosomal-ciliary protein implicated in autosomal recessive retinitis pigmentosa. *Adv Exp Med Biol*. 801:185-190.

Zamboni, L., and M. Stefanini. 1971. The fine structure of the neck of mammalian spermatozoa. *Anat Rec*. 169:155-172.

Zenker, J., M.D. White, R.M. Templin, R.G. Parton, O. Thorn-Seshold, S. Bissiere, and N. Plachta. 2017. A microtubule-organizing center directing intracellular transport in the early mouse embryo. *Science*. 357:925-928.
